# Nonparametric expression analysis using inferential replicate counts

**DOI:** 10.1101/561084

**Authors:** Anqi Zhu, Avi Srivastava, Joseph G. Ibrahim, Rob Patro, Michael I. Love

## Abstract

A primary challenge in the analysis of RNA-seq data is to identify differentially expressed genes or transcripts while controlling for technical biases. Ideally, a statistical testing procedure should incorporate the inherent uncertainty of the abundance estimates arising from the quantification step. Most popular methods for RNA-seq differential expression analysis fit a parametric model to the counts for each gene or transcript, and a subset of methods can incorporate uncertainty. Previous work has shown that nonparametric models for RNA-seq differential expression may have better control of the false discovery rate, and adapt well to new data types without requiring reformulation of a parametric model. Existing nonparametric models do not take into account inferential uncertainty, leading to an inflated false discovery rate, in particular at the transcript level. We propose a nonparametric model for differential expression analysis using inferential replicate counts, extending the existing SAMseq method to account for inferential uncertainty. We compare our method, Swish, with popular differential expression analysis methods. Swish has improved control of the false discovery rate, in particular for transcripts with high inferential uncertainty. We apply Swish to a single-cell RNA-seq dataset, assessing differential expression between sub-populations of cells, and compare its performance to the Wilcoxon test.

## Introduction

Quantification uncertainty in RNA-seq arises from fragments that are consistent with expression of one or more transcripts within a gene, or fragments which map to multiple genes. Abundance estimation algorithms probabilistically assign fragments to transcripts, and may simultaneously estimate technical bias parameters, providing a point estimate of transcript abundance. Transcript abundance can be compared across samples directly, or transcript abundances can be summarized within a gene to provide gene-level comparisons across samples. However, incorporating the uncertainty of the quantification step into statistical testing of abundance differences across samples is challenging, but nevertheless critical, as the inferential uncertainty is not equal across transcripts or genes, but is higher for genes with many similar transcripts (transcript-level uncertainty) or genes that belong to large families with high level of sequence homology (gene-level uncertainty).

Methods designed for transcript-level differential expression (DE) analysis include *BitSeq* [1], *EBSeq* [2], *Cuffdiff2* [3], *IsoDE* [4], and *sleuth* [5]. *BitSeq* uses MCMC samples from the posterior distribution during abundance estimation of each sample, to build a model of technical noise when comparing abundance across samples. A parametric Bayesian model is specified with the distribution for the abundance in a single sample being conjugate, such that the posterior inference comparing across samples is exact. *EBSeq* takes into account the number of transcripts within a gene when performing dispersion estimation, while *Cuffdiff2* models biological variability across replicates through over-dispersion, as well as the uncertainty of assigning counts to transcripts within a gene, using the beta negative binomial mixture distribution. *IsoDE* and *sleuth* leverage a measurement of quantification uncertainty from bootstrapping reads when performing DE analysis. These methods therefore may offer better control of FDR for transcripts or genes with high uncertainty of the abundance. *BitSeq, EBSeq, Cuffdiff2*, and *sleuth* incorporate biological variance and inferential uncertainty into a parametric model of gene or transcript expression, while *IsoDE* focuses on comparisons of bootstrap distributions for two samples at a time, aggregating over random pairs or all pairs of samples in two groups in the case of biological replicates. Another method, *RATs* [6], provides a test for differential transcript usage (DTU). *RATs* takes inferential uncertainty into account by repeating the statistical testing procedure across inferential replicates (posterior samples or bootstrap replicates), and then offers a filter on the fraction of replicate analyses which achieved statistical significance. In this study, we focus on differential transcript expression (DTE) and differential gene expression (DGE), the latter defined as a change in the total expression of a gene across all of its transcripts.

*BitSeq, EBSeq*, *Cuffdiff2*, and *sleuth* rely on parametric models of the transcript or gene abundance, assuming underlying distributions for the data and sometimes also for the parameters in the context of a Bayesian model. The distributions are typically chosen for being robust across many experiments and for efficient inference. In a recent simulation benchmark of DTE and DGE, *EBSeq* and *sleuth* did have improved performance for detecting DTE compared to methods designed for gene-level analysis, for *EBSeq* when sample size was large (*n* ≥ 6) and for *sleuth* when the sample size was low to moderate (*n* ∈ [3, 6]) [7], lending support for these models specialized for transcript-level inference. In this simulation benchmark, *EBSeq* and *sleuth* performed similarly to methods designed for detecting DGE, where uncertainty in abundance estimation is greatly reduced as described by Soneson et al. [8].

Previous studies have shown that nonparametric methods for differential expression analysis may be more robust compared to parametric models when the data for certain genes or transcripts deviates from distributional assumptions of the model [9]. Li and Tibshirani [9] proposed *SAMseq*, which has high sensitivity and good control of FDR at the gene-level, when the sample size is 4 per group or higher [7, 10, 11]. *SAMseq* uses a multiple re-sampling procedure to account for differences in library size across samples, and then averages a rank-based test statistic over the re-sampled datasets to produce a single score per gene. We considered whether the *SAMseq* model could be extended to account for inferential uncertainty during abundance estimation, as well as batch effects and sample pairing.

Further, we investigated whether a nonparametric method for comparison of counts, which accounts for inferential uncertainty, would provide benefit for new types of data exhibiting different distributions compared to bulk RNA-seq experiments. Single cell RNA-seq (scRNA-seq) provides researchers with high-dimensional transcriptomic profiles of single cells [12], revealing inter-cell heterogeneity and in particular sub-populations of cells that may be obscured in bulk datasets. Due to the limited amount of RNA material in a single cell and the low capture efficiency, all current protocols for scRNA-seq experiment use amplification techniques. To reduce the amplification bias, it is common to use Unique Molecular Identifiers (UMIs), short random sequences that are attached to the cDNA molecules per cell [13]. Ideally, by counting the total number of distinct UMIs per gene and per cell, one obtains a direct count of cDNA molecules, reducing bias from amplification.

Among the multiple different types of assays available for UMI-tagged scRNA-seq, droplet-based protocols have been of particular importance because of their relatively low cost, higher capture rate, robustness and accuracy in gene-count estimation. However, current dscRNA-seq (droplet-based single-cell RNA sequencing) protocols are limited in their resolution, as they only sequence the 3’ end of the mRNA molecule. This complicates full transcript-level quantification, and in fact such quantification is not possible for collections of transcripts that are not disambiguated within a sequenced fragment’s length of the 3’ end. Moreover, since dscRNA-seq protocols typically rely on considerable amplification of genomic material, the UMI sequences designed to help identify duplicate fragments are often subject to errors during amplification and sequencing. Most dscRNA-seq quantification pipelines [14–17] thus include some method for de-duplicating similar UMIs (i.e. UMIs within some small edit distance) attached to reads that align to the same gene. These dscRNA-seq quantification pipelines [14–17] simplify the problem of gene-level count estimation by discarding the gene-ambiguous reads, which represent a considerable portion of the data (12-23% as reported by Srivastava et al. [18]). *Alevin* [18] is the first tool to propose a method to account for sequencing and amplification errors in cellular barcodes (CB) and UMIs while also taking into account gene multi-mapping reads during quantification. To resolve gene ambiguity, *Alevin* first uses the Parsimonious UMI Graph (PUG) resolution algorithm. In the case that parsimony fails to distinguish a single best gene, abundance is estimated by utilizing the proportion of uniquely-mapping reads per gene to distribute ambiguous counts by an expectation-maximization procedure. *Alevin* also can produce inferential replicates in the form of multiple matrices of estimated counts, via bootstrapping of the counts in their PUG model. We assessed how incorporation of this inferential uncertainty using *Swish* resulted in different sets of DE genes compared to those called by a rank-based test ignoring uncertainty, in the context of calling DE genes between sub-populations of cells detected in scRNA-seq.

We describe a nonparametric method for gene- or transcript-level differential expression analysis, “SAMseq With Inferential Samples Helps”, or *Swish*, that propagates quantification uncertainty from Gibbs posterior samples generated by the *Salmon* [19] method for transcript quantification. In addition to the incorporation of uncertainty in the counts, *Swish* also extends the two group comparison in *SAMseq* by handling experiments with discrete batches or sample pairing. We show through simulation studies that the proposed method has better control of FDR and better sensitivity for DTE, compared to existing methods designed for transcript-level or gene-level analysis. For DGE, *Swish* tends to perform as well as existing methods designed for gene-level analysis, with better control of FDR than *SAMseq* for genes with high inferential uncertainty. We also compare *Swish*, leveraging estimation uncertainty, with a Wilcoxon test on gene expression in scRNA-seq sub-populations of cells in the developing mouse brain. *Swish* is available within the `fishpond` R/Bioconductor [20] package: https://bioconductor.org/packages/fishpond.

## 1 Materials and Methods

### 1.1 Median-Ratio Scaled Counts

In order to apply nonparametric rank-based methods to estimated counts per gene or transcript, we first remove biases on the counts due to library size differences or changes in the effective length of the gene or transcript across samples [19]. For 3’-tagged datasets, where we do not expect counts to be proportional to length, we correct only for library size but not for effective length, which is not computed. *scaledTPM* counts are generated which are roughly on the scale of original estimated counts, but are adjusted to account for technical biases with respect to effective transcript or gene length [8]. We perform median-ratio normalization

[21] of the *scaledTPM*, to remove any residual differences in *scaledTPM* across samples due to library size differences. The resulting scaled counts can then be directly compared across samples.

Suppose a matrix *Y*^0^ of estimated counts from *Salmon*. The rows of the matrix represents genes or transcripts, (*g* = 1,…,*G*), and columns represent samples, (*i* = 1,…,*m*). Let 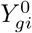 denote the count of RNA-seq fragments assigned to gene *g* in sample *i*. The estimated counts 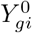 are divided by a bias term accounting for the sample-specific effective length of the gene or transcript, denoted as *l_gi_* [8, 22]. We first scale the *l_gi_* by the geometric mean over *i*, giving a bias correction term *b_gi_* centered around 1:

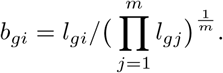

Then we divide the *b_gi_* from the estimated counts:

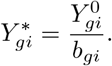

Again, for 3’-tagged data, we set *l_gi_* = *b_gi_* = 1 in the above. The 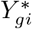 are scaled to the geometric mean of sequencing depth, by

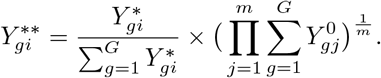

For each sample *i* a median-ratio size factor is computed as

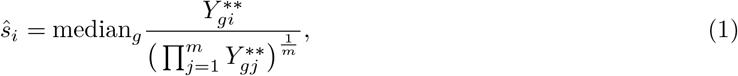

where the median is taken over *g* where the numerator and denominator are both greater than zero. For scRNA-seq data, an alternative size factor estimate is used, as described in a later section. Lastly, the scaled count used for nonparametric methods is given by:

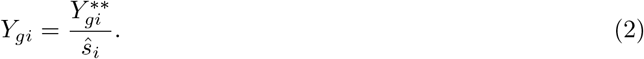

After scaling, we filter features (genes or transcripts) with low scaled counts and therefore low power: for bulk RNA-seq, we recommend keeping features with an estimated count of at least 10 across a minimal number of samples, and here we set the minimal number of samples to 3. For UMI de-duplicated scRNA-seq data, we lowered the minimum count to 3 and require at least 5 cells, as described in a later section.

We scale all of *Salmon*’s Gibbs sampled estimated count matrices using the above procedure. For posterior sampling, *Salmon* adopts a similar approach to *MMSEQ* [23] to perform Gibbs sampling on abundance estimates, alternating between (1) Multinomial sampling of fragment allocation to equivalence classes of transcripts, and (2) conditioning on these fragment allocations, Gamma sampling of the transcript abundance, as described in further detail in the Supplementary Note 2 of Patro et al. [19]. The Gibbs samples are thinned every 16th iteration to reduce auto-correlation (*Salmon*’s default setting), and 20 thinned Gibbs samples are used as the posterior samples in the following methods.

### 1.2 Inferential Relative Variance

For each gene (or transcript) from each sample, *S* Gibbs sampled estimated counts are generated using *Salmon*. Estimated counts are then scaled and filtered as described above.

We define the *inferential relative variance* (InfRV), per gene and per sample, as

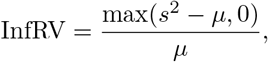

where *s*^2^ is the sample variance of scaled counts over Gibbs samples, and *μ* is the mean scaled count over Gibbs samples. In practice, we also add a pseudocount of 5 to the denominator and add 0.01 overall, in order to stabilize the statistic and make it strictly positive for log transformation. InfRV is therefore a measure of the uncertainty of the estimated counts, which is roughly stabilized with respect to *μ*. The larger the InfRV, the higher the uncertainty in quantification for that gene or transcript in that sample relative to other genes with the same mean. Note that InfRV is not used in the *Swish* statistical testing procedure, but only used in this work for visualization purpose, to categorize genes or transcripts by their inferential uncertainty, in a way that is roughly independent of the range of counts for the feature.

### 1.3 Test Statistic over Gibbs Samples

In *Swish*, we consider experiments with samples across two conditions of interest. We additionally allow for two or more batches to be adjusted for, or comparisons of paired data, described below. We first consider an experiment with two conditions and no batches. Let *m_k_* denote the sample size in condition *k, k* = 1, 2, and *m*_1_ + *m*_2_ = *m*. Let *C_k_* denote the collection of samples that are from condition *k*.

For the following, consider a single matrix 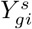 of scaled counts from one of the Gibbs samples, (*s* = 1,…, *S*). For gene *g*, the scaled counts 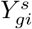 estimated from (2) are ranked over *i*. To break ties in 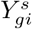 over the *m* biological samples, a small random value is draw from Uniform(0, 0.1) and added before computing the ranks, as in Li and Tibshirani [9]. Let 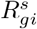 denote the rank of sample *i* among *m* biological samples for gene *g*. As in Li and Tibshirani [9], for a two group comparison we use the Mann-Whitney Wilcoxon test statistic [24] for gene *g*:

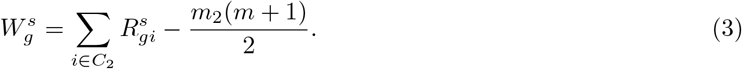

Under the null hypothesis that gene *g* is not differentially expressed, 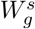 has expectation zero.

For every gene *g* = 1,…,*G*, we calculate 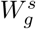 for all Gibbs sampled scaled count matrices (*s* = 1,…,*S*). As in Li and Tibshirani [9], we compute the mean over imputed count matrices as the final test statistic for gene *g*:

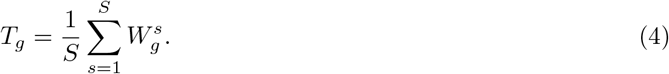

The main difference between *Swish* and *SAMseq* is the use of inferential replicate count matrices from an upstream quantification method as input to the Mann-Whitney Wilcoxon test, as opposed to Poisson down-sampled count matrices. Additionally, *Swish* extends *SAMseq* for various other experimental designs, described below.

### 1.4 Stratified Test Statistic

In experiments with two or more batches, the above test statistic can be generalized to a stratified test statistic as described in Van Elteren [25]. We use the following to produce a summarized test statistic:

1. Stratify the samples by the nuisance covariate (e.g. batch). Suppose we have *J* strata. In stratum *j* the sample size for condition *k* is *m_j,k_* and the total sample size is *m_j_*, i.e. *m*_*j*,1_ + *m*_*j*,2_ = *m_j_*.
2. For stratum *j*, compute the test statistic 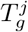.
3. The final test statistic is calculated as the weighted combination of 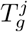 over *J* strata

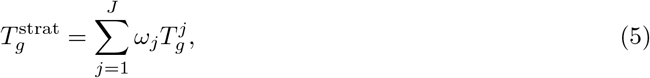

where 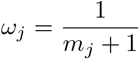 is the weight assigned to strata *j*.

When the difference in expression for gene *g* is consistent across batches, the weighted test statistic maximizes the power of test [26].

### 1.5 Signed-Rank Statistic for Paired Data

In experiments with paired samples, e.g. before and after treatment, we use a Wilcoxon signed-rank test statistic [24] in (4) in lieu of the Mann-Whitney Wilcoxon statistic (3). For pair index *p* = 1,…,*m*/2:

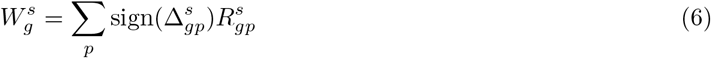

where 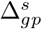 is the difference in 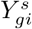 for the two samples in pair *p*, and 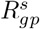 is the rank over pairs of 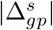.

Unlike the Mann-Whitney Wilcoxon and stratified version described previously, the signed-rank statistic is not necessarily identical after transformation of the data by a monotonic function, e.g. by the square root or shifted logarithm. We choose to keep the data as scaled counts when computing the signed-rank statistic, and not to perform the square root or other transformation, so as to promote the association of higher ranks to matched samples with higher scaled counts for a given gene or transcript.

### 1.6 Mann-Whitney Wilcoxon for Differences across Two Groups

In experiments with two groups of paired samples, e.g. before and after treatment, with pairs of samples divided across two treatments, one may be interested in testing if the effect of treatment is the same across the two groups. Here, we use a Mann-Whitney Wilcoxon test statistic on the log fold changes computed on the pairs, again taking the mean of this statistic over inferential replicates. This test will reject the null hypothesis for genes or transcripts where the multiplicative effect on counts differs across the two treatments, while controlling for any differences at baseline. It is analogous to testing an interaction term in linear model. The log fold change is chosen here, as opposed to the signed-rank statistic chosen above, as the signed-rank statistic would be sensitive to differences of the baseline counts across the two treatments, which is not desired.

### 1.7 Permutation for Estimation of FDR

Under the null hypothesis that gene *g* is not differentially expressed, the distribution of test statistic *T_g_* in (4)-(6) is not known. Instead, following the approach in *SAMseq* [9] we use permutation tests to construct the null distribution for *T_g_*. In the case of stratified data, we perform permutations independently within strata and calculate a weighted test statistic as in (5). For paired data, we randomly permute condition labels within pairs. Finally we use the permutation plug-in method [27] to estimate the q-value for each gene *g*, leveraging functions in the R package *qvalue* [27]. We provide the option to estimate π_0_, the proportion of nulls, from the observed test statistics and the permutation null distribution, but by default we set 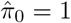, which reduces to the method of Benjamini-Hochberg [28] for FDR control. In practice, 4 samples per group is sufficient for the permutation method to have sensitivity to detect differential expression [9, 11].

### 1.8 Simulation of Bulk RNA-seq

In order to evaluate *Swish* and other methods, we make use of a previously published simulated RNA-seq dataset [7], which we briefly describe here. *polyester* [29] was used to simulate paired-end RNA-seq reads across four groups of samples: two condition groups with various differential expression patterns described below, which were balanced across two batches. One batch demonstrated realistic fragment GC bias, and one batch demonstrated approximately uniform coverage. The fragment GC bias was derived via *alpine* [30] from GEUVADIS samples [31]. Likewise, the baseline transcript expression was derived from a GEUVADIS sample. Each condition group had 12 samples in total, equally split among the two batches, giving 24 samples in all. Fragments were simulated from 46,933 expressed transcripts belonging to 15,017 genes (Gencode [32] release 28 human reference transcripts).

70% of the genes were simulated with no differential expression across condition, while 10% of the genes had all of their transcripts differentially expressed with the same fold change, 10% of the genes had a single transcript differentially expressed, and 10% of the genes had shifts in expression among two of their transcript isoforms which resulted in no change in total expression. For the purpose of gene-level differential expression, 20% of the genes were non-null due to changes in total expression. For transcript-level differential expression, all transcripts expressed in the first category were differentially expressed, as well as one transcript per gene from the second category, and two transcripts per gene from the third category.

Transcripts were quantified using *Salmon* v0.11.3 and *kallisto* [33] v0.44.0, with bias correction enabled for both methods, and generating 20 Gibbs samples for *Salmon* and 20 bootstrap samples for *kallisto*. The scripts used to generate the simulation are provided in the Code Availability section, and links to the raw reads and quantification files are listed in the Data Availability section.

### 1.9 Simulation Evaluation

We evaluated methods across four settings:

1. DGE analysis with one batch of samples (fragment GC bias), 6 samples in each of the two conditions,
2. DGE analysis with two balanced batches of samples, 12 samples in each of the two conditions,
3. DTE analysis with one batch of samples (fragment GC bias), 6 samples in each of the two conditions,
4. DTE analysis with two balanced batches of samples, 12 samples in each of the two conditions.

To evaluate the performance of methods on DGE and DTE, we use the iCOBRA package [34] to construct plots of the true positive rate (TPR) over the false discovery rate (FDR) at three nominal FDR thresholds: 1%, 5%, and 10%. The methods compared include *Swish, DESeq2* [11], *EBSeq* [2], *limma* with *voom* transformation [35], *SAMseq* [9], and *sleuth* [5]. We chose a set a methods to represent popular choices among classes of approaches. We note that *edgeR* [36] and *edgeR* with quasi-likelihood [37] performed well in a previous benchmark of this dataset [7], with both performing similarly to *limma*. *EBSeq* and *sleuth* are designed for DTE or DGE analysis taking the inferential uncertainty of quantification into account, while *SAMseq*, *DESeq2*, and *limma* are designed for gene-level analysis, but have been shown nevertheless to be able to recover DTE when supplied with transcript-level estimated counts [7]. We provided minimal filtering across all methods, removing transcript or genes without an estimated count of 10 or more across at least 3 samples. We also used the default recommended filtering functions `filterByExpr` and `sleuth_prep` for *limma* and *sleuth*, respectively.

For *DESeq2*, *EBSeq*, *limma*, and *SAMseq*, we provided the methods with counts on the gene- or transcript-level generated by the *tximport* package [8], using the *lengthScaledTPM* method. For the two balanced batches experiment, we provided all methods with the batch variable, except *SAMseq* which does not have such an option. For *SAMseq*, for the two batch experiment, we first adjusted the counts using the known batch variable and the `removeBatchEffect` function in the *limma* [38] package, which improved *SAMseq*’s performance. For *sleuth* we directly provided *kallisto* quantification files to the method, and for detection of DGE, changes in the total expression of all the transcripts of a gene across condition, we used the count summarization option.

Nonparametric methods have the advantage of being robust to outliers in sequencing data. To evaluate the robustness of our method, we introduced outliers into the counts of three randomly chosen transcripts that were not differentially expressed in the simulation. For each of these transcripts, we randomly selected one sample among the 12 total samples, and multiplied the estimated counts and the Gibbs samples by 1,000. We compared the q-values or adjusted p-values of these three transcripts from *Swish*, *limma*, *DESeq2*, and *EBSeq* (*Swish*, compared to three parametric methods that can take as input the modified count matrices). For *DESeq2*, we turned off the Cook’s-distance-based filtering procedure, to demonstrate the sensitivity of the un-filtered test results to outliers.

### 1.10 Highly replicated RNA-seq Datasets

To assess the performance of *Swish* on real bulk RNA-seq datasets, in particular its control of the target FDR, we downloaded two datasets that contain a large number of biological replicates, which make them useful for benchmarking statistical methods for differential expression analysis. The first dataset includes 43 high quality biological replicates of wild type *Saccharomyces cerevisiae* [39]. Random sets of 10, 20, 30, and 40 samples were chosen from the total of 43 samples, and randomly split into two groups to form test datasets. For each sample size, we repeat the resampling process 100 times. The second dataset includes 16 high quality biological replicates of wild type *Arabidopsis thaliana* [40]. Again, random sets of 10 samples, or the entire dataset of 16 samples was chosen, and randomly split into two groups. Partitions of the data were chosen to balance the number of samples from batch 1, as we found this batch to separate from batches 2 and 3 in exploratory data analysis (Supplementary Figure 1). Again for each sample size, we repeated the resampling process 100 times. For the yeast dataset, we focused on gene expression, while for the *Arabidopsis* dataset, we investigated both gene and transcript expression. As the samples in all cases were from the same experimental condition, and were chosen to balance with respect to technical variation when present, we do not expect any genes or transcripts to be reported as differentially expressed. We focused on *Swish*, *SAMseq*, and *limma* for this analysis, as *limma* had strong control of the FDR in many cases in the simulated data, and because *Swish* is an extension of the *SAMseq* method. For each method, on each random null dataset we recorded the proportion of genes or transcripts reported in an FDR set for a 5% target, which can be interpreted as a false positive rate (FPR).

### 1.11 Single Cell RNA-seq Simulation

In order to assess the ability of *Swish* to make use of uncertainty information contained in inferential replicates for single cell RNA-seq, we constructed a simulation combining the *splatter* [41] and *polyester* [29] Bioconductor packages. We used the human reference cDNA transcripts from Ensembl release 95, and simulated gene expression for 35,583 genes. Read counts were simulated for 40 cells in two groups using *splatter* with *DE factor location* of 3 on the log2 scale, and *DE factor scale* of 1 on the log2 scale. 10% of genes were chosen to be differentially expressed. Stranded single-end reads of length 100 bp were generated using *polyester*, drawing from the 400 bp closest to the 3’ end of one of the transcripts per gene, unless the transcript was shorter than 400 bp in which fragments were drawn from the entire transcript. This aspect of the simulation was chosen to best reflect 3’-tagged scRNA-seq data, in which there is more uncertainty in assigning reads to genes, compared to full-length RNA-seq data. Gene-level expression was estimated by running *Salmon* using the Ensembl full length cDNA reference transcripts as the index, without length correction (as recommended for 3’-tagged reads) and summarizing counts to the gene level with *tximport*.

### 1.12 Single Cell RNA-seq of Mouse Neurons

We downloaded a dataset of scRNA-seq of the developing mouse brain in order to evaluate the performance of *Swish* on RNA-seq counts with a different distribution and with more gene-level inferential uncertainty than bulk RNA-seq. In this dataset, cells from the cortex, hippocampus and ventricular zone of a embryonic (E18) mouse were dissociated and sequenced on the 10x chromium pipeline. scRNA-seq data was quantified by *Alevin* [18] v0.13.0 with the default command line parameters along with `--numCellBootstraps 30` to generate 30 bootstrap estimated count matrices. The total number of cells was 931, and the number of genes was 52,325 (Gencode [32] release M16 mouse reference transcripts).

In a comprehensive benchmark, Soneson and Robinson [42] compared DE methods designed for both scRNA-seq and bulk RNA-seq, and found that the Mann-Whitney Wilcoxon test [24] had favorable performance compared to numerous other methods in detection of DE genes across cells. However, the Wilcoxon test was listed as having the downside of not accommodating complex designs (for example, covariate control, sample pairing, or interaction terms). We therefore compared *Swish* to the Wilcoxon test statistic in detection of genes differentially expressed across clusters.

*Seurat* [43] was used to cluster the cells in the dataset. First, cells with low quality were removed, based on quality control plots (Supplementary Figure 2). Those cells having less than 1,500 or greater than 6,500 genes expressed were removed based on the quality control plots. Additionally, those cells with greater than 4% percent of mitochondrial genes detected were removed. These filters resulted in 835 cells remaining in the dataset. For *Seurat* clustering, gene counts were then normalized by their total count, multiplied by 10,000, and the shifted logarithm was applied, with a shift of 1 to preserve zeros.

Only highly variable genes were used for clustering by *Seurat*, which was determined by calculating the mean expression of scaled counts and dispersion for each gene, binning genes by mean expression into 20 bins, and calculating z-scores for *s*^2^/*μ*, (the sample variance over the sample mean) within each bin. The highly variable genes were defined as those with mean expression between 0.0125 and 3 on the natural log scale and z-score of 0.5 or larger. A linear model was fit to the log scaled counts, to further remove any technical artifacts associated with the total number of UMIs detected per cell and with the percent of mitochondrial genes detected. The top 10 principal components of residuals of highly variable genes were used to cluster the cells, using *Seurat*’s graph-based unsupervised clustering method.

After defining clusters using the highly variable genes, we returned to the matrix of estimated counts and attempted to find genes which were differentially expressed across two specific clusters. The clusters were chosen based on expression of neural differentiation marker genes reported by a previous study [44]. We applied both *Swish* and a standard two-sample Wilcoxon test to the scaled counts of cells in the two clusters. We scaled the counts using size factors estimated with the setting `type=“poscounts”` in the *DESeq2* package [11], as recommended for scRNA-seq differential expression analysis [45]. Genes without a count of 3 or more in 5 or more cells were removed prior to the DE analysis across clusters. *Swish* produces a q-value, while the p-values from the Wilcoxon test were adjusted using the Benjamini-Hochberg [28] method to produce nominal FDR bounded sets.

To assess the performance of *Swish* compared to the two-sample Wilcoxon test on real scRNA-seq data, we sought an independent dataset that we could regard as a pseudo-gold-standard. As scRNA-seq may have its own systematic biases, we focused for validation on a bulk RNA-seq dataset also of embryonic (E14.5) mouse brain, which employed laser-capture microdissection in order to isolate various regions of the brain [46]. We downloaded and quantified 5 samples from the ventricular zone (VZ) and 5 samples of the cortical plate (CP) using *Salmon*, and then analyzed the bulk RNA-seq data on the gene-level using *tximport* followed by *DESeq2*.

## 2 Results

### 2.1 Simulation of Bulk RNA-seq

We evaluated the InfRV in the simulated bulk RNA-seq dataset at the gene and transcript level. At the transcript level, there are two main sources of abundance uncertainty: 1) among different transcripts within a gene, and 2) due to homologous sequence of transcripts across genes. At the gene level, we observed substantially less abundance uncertainty, as expected (Figure 1). At the gene level the uncertainty among transcripts within a gene has been collapsed, and only the latter source of uncertainty remains, also noted by Soneson et al. [8].

**Figure 1:**
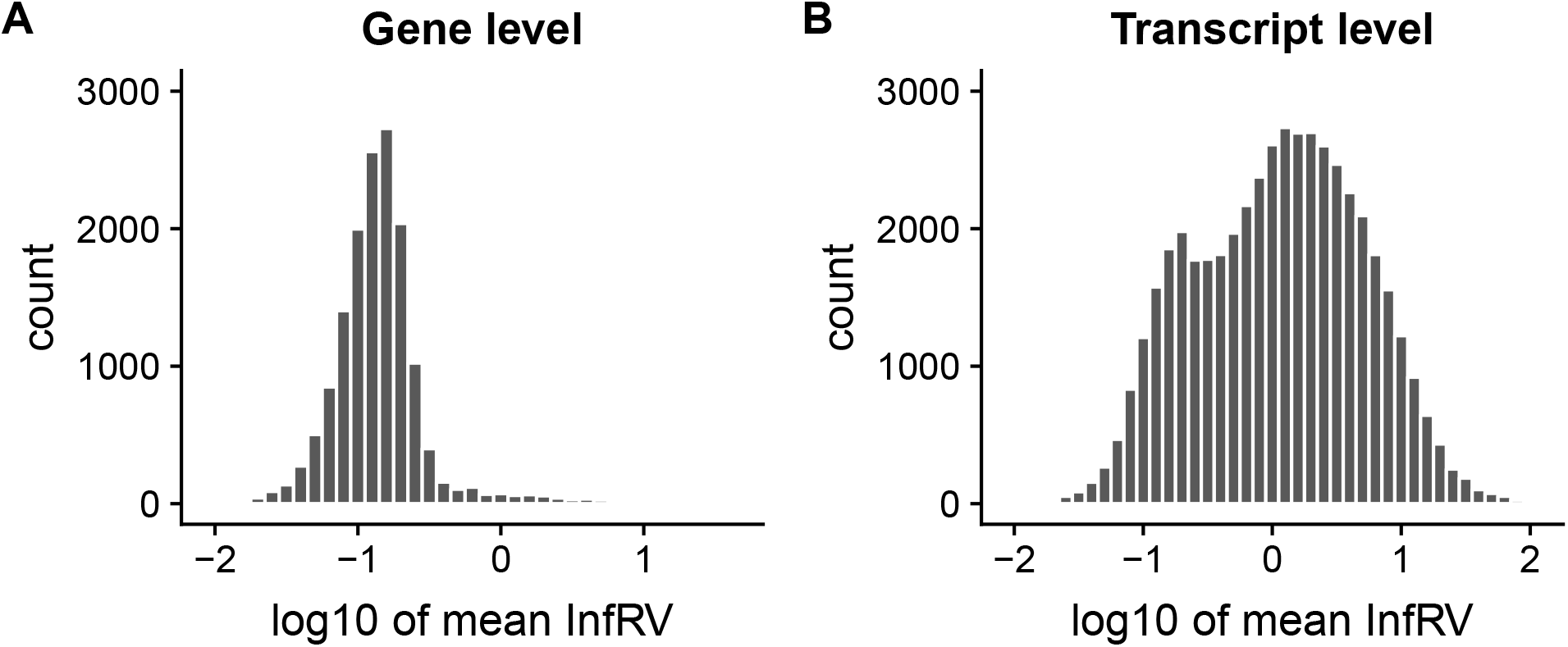
Mean of InfRV over samples for genes and transcripts in the simulation studies (log10 scale). InfRV is binned by increments of 0.1, so that 10 bins span an order of magnitude. (A) Gene level for the single batch experiment. (B) Transcript level for the single batch experiment.

At the gene level, where there was less quantification uncertainty, *Swish* had similar performance as DGE methods overall (Figure 2). For the experiment with a single batch, *Swish* had similar TPR and FDR to *EBSeq*, *limma*, *SAMseq*, and *sleuth*, and better control of FDR compared to *DESeq2*. *Swish* had improved FDR control compared to *SAMseq*, but only slightly so, and this minor performance difference was expected as there was relatively low InfRV compared to transcript-level analysis. For the experiment with two batches, *Swish*, *limma*, and *EBSeq* showed the best performance, with good sensitivity and control of FDR. *Swish* had tight control of FDR for the 5% and 10% target while *SAMseq* had observed FDR of 10% for the 5% target, and observed FDR of 18% for the 10% target (Supplementary Figure 3). *sleuth* exhibited loss of control of the FDR for the 5% and 10% target in the two batch experiment (Supplementary Figure 4).

At the transcript level, *Swish* had consistently high sensitivity and good control of FDR, where other methods either had low sensitivity or loss of FDR control in some setting. *Swish* controlled FDR for all nominal FDR targets (1%, 5%, and 10%), while *SAMseq* tended to exceed the 5% and 10% target for transcripts with medium-to-high InfRV, both for the single batch experiment (Figure 3), and the two batch experiment (Supplementary Figure 5). As opposed to gene-level analysis where there is relatively low InfRV, at the transcript level the performance difference between *Swish* and *SAMseq* is most obvious for the transcripts with the most quantification uncertainty, as expected. This can be seen in the bottom right panels of Figure 3 and Supplementary Figure 5, which show the top third of transcripts in the dataset by InfRV. *limma* had reduced sensitivity for all InfRV categories, for both experiments, due to filtering of DTE transcripts by `filterByExpr` [47]. With a less stringent filtering rule, *limma* showed improved sensitivity compared to its default settings, but still more than 10% below the sensitivity of other methods controlling the target FDR including *Swish* (Supplementary Figure 7). *EBSeq* had high sensitivity but loss of FDR control for the transcripts with high InfRV. *sleuth* performed well in the single batch experiment, but lost control of FDR for the 5% and 10% target in the two batch experiment (Supplementary Figure 6).

**Figure 2:**
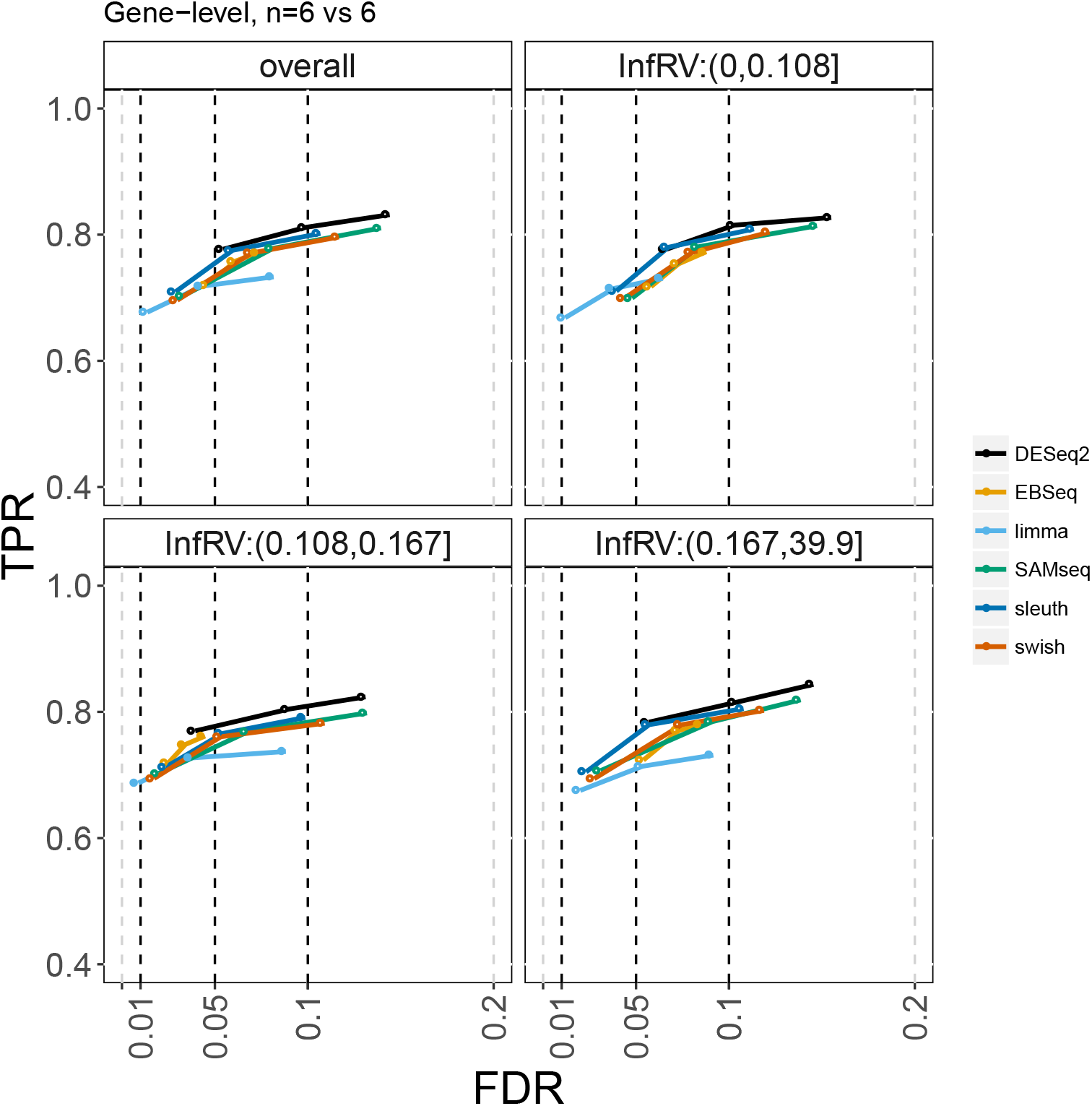
True positive rate (y-axis) over false discovery rate (x-axis) for DGE analysis with only one batch of samples, with 6 samples in each of the two conditions. The first panel depicts overall performance, while the subsequent three panels depict the genes stratified into thirds by InfRV averaged over samples.

We assessed the Mann-Whitney Wilcoxon test on scaled counts, with linear-model-based removal of batch effect for the two batch experiment, followed by multiple test correction with the method of Benjamini-Hochberg [28] to control FDR. In the gene-level and transcript-level experiments, for both one and two batch experiments, use of the Mann-Whitney Wilcoxon test performed similarly to *SAMseq* (Supplementary Figures 8-11).

We examined how the methods performed on the simulation across a finer grid of sample sizes. We focused on transcript-level analysis, and on the one batch experiment, and varied the per-group sample size over the range, *n* ∈ {3, 4, 5, 9, 12, 15, 18, 20}. At sample sizes 3 and 4, *Swish* returned the same set of transcripts for the 1%, 5%, and 10% target FDR, which controlled at 10% FDR for n=3, and controlled at 1%, 5% and 10% for n=4 (Supplementary Figures 12 and 13). In contrast, *SAMseq* returned no transcripts for any target FDR at n=3, and no transcripts for the 1% target FDR at n=4. The n=5 experiment (Supplementary Figure 14) had similar results to n=6 (Figure 3), but with lower sensitivity for *SAMseq* for the 1% target. In the experiments from n=9 to n=20, the top third transcripts by InfRV revealed a difference between *Swish* and *SAMseq*, where the latter exceeds the 5% and 10% FDR targets. In these higher sample size experiments, *sleuth* did not control FDR for the 5% and 10% target, or the 1% target for *n* starting at 15 (Supplementary Figures 20-24).

**Figure 3:**
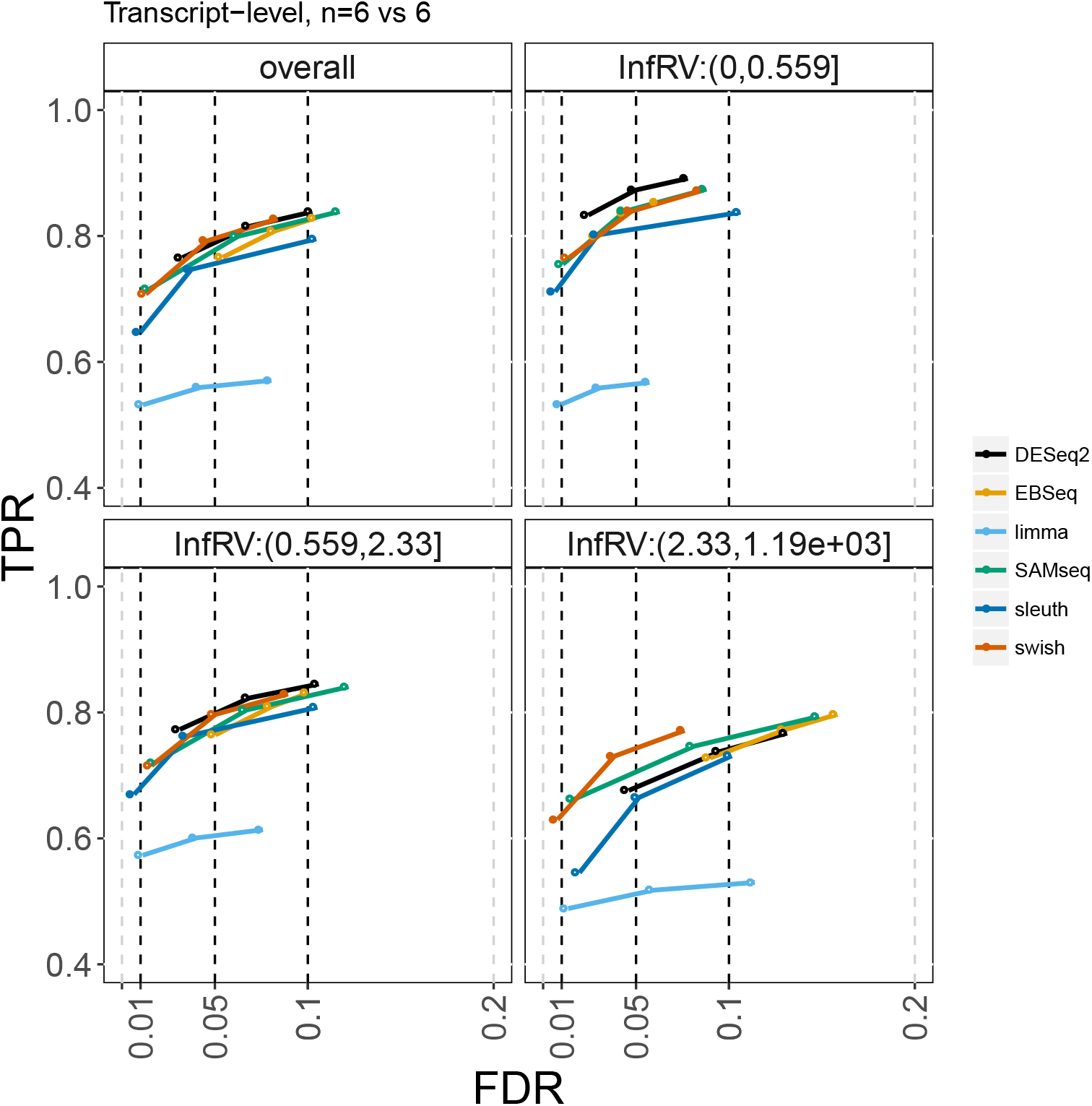
True positive rate (y-axis) over false discovery rate (x-axis) for DTE analysis with only one batch of samples, with 6 samples in each of the two conditions. As in Figure 2, the first panel depicts overall performance, while the subsequent three panels depict the transcripts stratified into thirds by InfRV averaged over samples.

When the count for a single sample for a null transcript was modified to be an extreme outlier, *Swish* did not call this transcript as differentially expressed, with q-value close to one, as did *limma* with its adjusted p-values (Supplementary Table 1). *DESeq2* with its outlier flagging procedure turned off and *EBSeq* with default settings returned very small adjusted p-values for these genes with inserted outliers. This case demonstrated another benefit of nonparametric models in their robustness to model mis-specification or outliers, also demonstrated by Li and Tibshirani [9].

Overall, *Swish* performed well at the gene level, out-performing the method *SAMseq*, which it is based upon, in the two batch experiment in terms of FDR control. *Swish* out-performed other methods at the transcript level, both for the single and two batch experiments, where other methods either had reduced sensitivity or loss of FDR in some setting.

### 2.2 Highly replicated RNA-seq Datasets

In the two highly replicated bulk RNA-seq datasets *Swish* had lower false positive rate compared to *SAMseq*, and similar to *limma*. In both datasets, in which no features should be detected as differentially expressed by construction, *Swish* in particular had fewer random partitions in which more than 5% of the features were called differentially expressed. In the yeast dataset *Swish* rarely called more than 5% of the features as differentially expressed, while *SAMseq* had more such iterations where more than 5% or even up to 40% of the features would be called differentially expressed (Supplementary Figure 25-28). Overall *limma* tended to call very few genes as differentially expressed in the null comparisons. Similar relative performance could be seen in the *Arabidopsis* dataset, though overall there were fewer iterations with high rates of false positive calls for any method, compared to the yeast dataset. At the transcript level, there were rarely any false positive calls for any method (Supplementary Figure 29 and 30). At the gene level, there were still few calls, but *SAMseq* tended to have more iterations with nonzero false positive rate, compared to the other two methods (Supplementary Figure 31 and 32).

### 2.3 scRNA-seq Simulation

The global performance of *Swish* and the Wilcoxon test was similar on the *splatter-polyester* simulation. Still, we identified a number of individual null genes in which the Wilcoxon test followed by Benjamini-Hochberg adjustment resulted in a very low adjusted p-value, while *Swish* did not detect these genes at a 5% nominal FDR target. After simulating scRNA-seq counts for two groups of 20 cells each, and testing at a target 5% FDR, *Swish* had 91.7% sensitivity and an observed FDR of 5.3%, while the Wilcoxon test had 92.4% sensitivity and an observed FDR of 5.7%. However, we detected a number of null genes with spurious expression values in an MA-plot of the dataset (Supplementary Figure 33). These null genes were assigned non-zero estimates of expression — and differentially so across the two simulated groups of cells — due to sequence homology with other genes which were simulated as differentially expressed. We focused on 14 genes that could be identified in this MA plot, which had a total count across cells > 5, had a t-statistic > 5, and had InfRV > .2. For all of these 14 genes, *Swish* gave a greatly increased q-value (less significant) compared to the adjusted p-value from the Wilcoxon test (Supplementary Figure 34), such that 9 of the genes were not detected in a 5% nominal FDR set. Plotting the inferential replicates for all of the 40 cells for these genes revealed that there was extensive inferential uncertainty for the non-zero counts for these genes (Supplementary Figure 35). While examining only a point estimate of the expression may lead a method to infer differential expression for such null genes, taking the inferential uncertainty into account during the statistical testing lead to more correct results for these genes, while not affecting the overall sensitivity of the method for truly differentially expressed genes.

### 2.4 scRNA-seq of Mouse Neurons

We evaluated the extent to which inclusion of uncertainty in the gene-level abundance would affect differential expression in an scRNA-seq dataset. As described in the Methods, *Seurat* was used to identify clusters in a dataset of 835 cells from embryonic mouse brain. *Seurat* identified a total of nine clusters, visualized in Supplementary Figure 36. Expression level of known cell-type markers for developing mouse brain, as reported by Loo et al. [44], was overlaid on the t-Distributed Stochastic Neighbor Embedding (tSNE) [48] plot. *Eomes*, a gene expressed in neural progenitor cells, was highly expressed in cluster 7, while *Neurod2*, a marker of differentiated neurons, was highly expressed cluster 5 (Supplementary Figure 37). We compared the test results of a two-sample Mann-Whitney Wilcoxon [24] test to *Swish* across groups of cells from cluster 5 and cluster 7. We defined sets of genes which were called by *Swish* or the Wilcoxon test at a 5% threshold.

We used an independent bulk RNA-seq dataset as a pseudo-gold-standard, to validate the calls of *Swish* and the Wilcoxon test [46]. This dataset contains bulk RNA-seq of 5 samples from the ventricular zone (VZ) and 5 samples from the cortical plate (CP) of mice embryos of a similar age (E14.5) to the mice embryos in the scRNA-seq data (E18). The brain regions were isolated using laser-capture microdissection. We chose the VZ and CP regions from the Fietz et al. [46] dataset to have highest correlation with clusters 7 and 5 from the scRNA-seq dataset: the VZ is enriched with progenitor cells marked by *Eomes*, while the CP is enriched with differentiated neurons marked by *Neurod2*. We refer to the bulk RNA-seq dataset as a “pseudo-gold-standard”, as the regions produced by microdissection contain various cell types, and so the comparison of bulk with *in silico* sorted single cells is not exact. Nevertheless, a Pearson correlation of 0.61 was obtained when comparing the log2 fold changes from the scRNA-seq, cluster 7 vs cluster 5, with the log2 fold changes from the bulk RNA-seq, VZ vs CP, so we felt the pseudo-gold-standard would therefore be useful for assessing method performance.

Comparing random subsets of 20 cells each (40 cells total) from cluster 7 and cluster 5 did not reveal large differences in performance between *Swish* and Wilcoxon on this particular dataset, in contrast to the clear enrichment of false positives seen in the *splatter-polyester* scRNA-seq simulation. However, when randomly selecting subsets of 10 cells each (20 cells total) from the two clusters, *Swish* had a clear advantage over Wilcoxon tests in terms of number of genes recovered at the same observed FDR level (Supplementary Figure 38). For the 5% and 10% target FDR, *Swish* was able to call 50 more genes than the Wilcoxon test, at comparable observed FDR, increasing the number of reported genes by more than half at 5% target FDR and by more than one third at 10% target FDR. Both *Swish* and Wilcoxon test were close to hitting their 5% and 10% FDR targets on the dataset of 20 cells total, while they both tended to exceed the targets on the dataset with 40 cells total. Some excess may be attributable to the fact that the VZ and CP brain regions comprise more than the individual cell populations isolated in clusters 7 and 5 from the scRNA-seq analysis.

## 3 Discussion

Many parametric models have been proposed to include quantification uncertainty into statistical testing procedures when evaluating differential transcript or gene expression. We extended an existing nonparametric framework, *SAMseq*, through the incorporation of inferential replicate count matrices, so that the test statistic better reflects the uncertainty of the counts. We show that this extension recovers control of FDR in particular for transcripts with high inferential relative variance. We used posterior samples generated by a Gibbs sampler with thinning to reduce autocorrelation, although bootstrapping of reads to generate inferential replicates is also compatible with our proposed method. *Swish* additionally extends *SAMseq* by allowing for control of discrete batches using the stratified Wilcoxon test, and allowing for analysis of paired samples using the Wilcoxon signed-rank test. We note that *Swish* makes use of inferential replicates that provide information about uncertainty due to the assignment of reads. Other sources of uncertainty, such as experimental uncertainty, may not be captured by the inferential replicates. A simple example is the uncertainty of expression for genes which receive no reads from a lowly sequenced sample. Nevertheless, previous work has shown that in some cases, correct quantification can be recovered despite missing fragments due to common technical biases in sequencing data, to the degree that the bias can be modeled [19, 30].

*Swish* has the greatest advantage over other competing methods at the transcript level, while it performs similar to other methods at the gene level. We additionally demonstrated the use of *Swish* on scRNA-seq counts generated by the *Alevin* method, making use of bootstrap inferential replicates at the gene level. On simulated scRNA-seq data, we found clear evidence of false positives which would be called by a standard rank-based method, which are appropriately not called as differentially expressed by *Swish* due to its reliance on inferential replicates in the testing procedure. On the mouse neurons dataset, we found that *Swish* outperformed the Wilcoxon test for comparisons of small numbers of cells (e.g. 20 cells total), but performed similar for larger sets of cells, at least in so far as we could detect using a pseudo-gold-standard dataset for validation.

We note that, although we assessed the method on a real dataset from the 10x chromium pipeline, *Alevin* followed by *Swish* on inferential replicates could likewise be performed on other platforms such as Fluidigm – though *Alevin*’s bias model could be further optimized for alternate platforms. We anticipate that taking into account uncertainty from inferential replicates for scRNA-seq will be equally valuable for gene-level differential expression for any 3’-tagged protocol, and for transcript-level differential expression for any full length protocol. In this scRNA-seq analysis, we did not take into account that we first clustered the data based on highly variable genes, and then performed differential analysis using those and other genes, which may generate anti-conservative inference as noted recently by Zhang et al. [49]. We are interested in extensions which may alleviate this problem and in general help to avoid comparisons across unstable clusters [50–53].

Some of the limitations of our proposed work include the types of experiments that it can be used to analyze. Because it relies on permutation of samples to generate a null distribution over all transcripts or genes, *SAMseq* and *Swish* in practice do not necessarily have sufficient power to detect differential expression when the sample size is less than 4 per group. Additionally, certain terms which can be included in linear models, such as controlling for continuous variables, cannot be easily incorporated into the nonparametric framework. We focused on inference across a discrete nuisance covariate, for example sample preparation batches, which motivated the stratified test statistic, as well as inference across a discrete secondary covariate of interest, for example a secondary treatment, which motivated the difference-across-two-groups method. However, we cannot yet accommodate more continuous measures of batch, such as those estimated by *RUVSeq* [54] or *svaseq* [55], unless such factors of unwanted variation could be discretized without substantial loss of information. Currently, *Swish* supports two group comparisons, with discrete covariate correction, sample pairing, and comparison of condition effects across a secondary covariate, although we could extend to other analysis types already supported by *SAMseq*, including multi-group comparisons via the Kruskal-Wallis statistic, quantitative association via the Spearman correlation, or survival analysis via the Cox proportional hazards model.

## 4 Code Availability

The `swish` function is available in the *fishpond* R/Bioconductor [20] package at https://bioconductor.org/packages/fishpond. The implementation is efficient, with vectorized code for row-wise ranking calculation via the *matrixStats* package [56]. The software runs in comparable or faster time to *DESeq2*, taking 1-2 seconds per 1,000 genes for small datasets, e.g. total *n* ∈ [10 – 20]. For larger datasets (total *n* > 100), *Swish* scales better than *DESeq2* (Supplementary Figure 39).

The package contains a detailed vignette with live code examples, showing how to import *Salmon* quantification files using the *tximeta* package, perform differential transcript or gene expression analysis with `swish`, and visualize results for each transcript or gene. The package vignette includes an example of a paired RNA-seq analysis, both a two group example and a differences across groups example. The software vignette leverages a subset of the RNA-seq data from a recent publication on human macrophage immune response [57] (Supplementary Figure 40).

The scripts used to generate the Love et al. [7] simulation data are available at the following link: https://dx.doi.org/10.5281/zenodo.1410443. The scripts used in evaluating the methods here, and in analyzing the scRNA-seq dataset have been posted at the following link: https://github.com/azhu513/swishPaper.

The versions of software used in the paper are: *fishpond* (*Swish* method): 0.99.32, *samr* (*SAMseq* method): 2.0, *qvalue*: 2.14.1, *Salmon*: 0.11.3, *Alevin*: 0.13.0, *tximport*: 1.10.1, *kallisto*: 0.44.0, *sleuth*: 0.30.0, *DESeq2*: 1.22.2, *EBSeq*: 1.22.1, *limma*: 3.38.3, *Seurat*: 2.3.4

## 5 Data Availability

The simulated data used to evaluate the methods is published in Love et al. [7]. The raw read files can be accessed at the following links: https://doi.org/10.5281/zenodo.1291375, https://doi.org/10.5281/zenodo.1291404, https://doi.org/10.5281/zenodo.2564176, https://doi.org/10.5281/zenodo.2564261. The processed simulation quantification files can be accessed at the following link: https://doi.org/10.5281/zenodo.2564115. The mouse neurons scRNA-seq dataset is available under the heading *Chromium Demonstration (v2 Chemistry) > Cell Ranger 2.1.0 > 1k Brain Cells from an E18 Mouse*, from https://support.10xgenomics.com/single-cell-gene-expression/datasets. The mouse brain isolated bulk tissue RNA-seq data are available from the Gene Expression Omnibus with the accession number GSE38805.

## 6 Acknowledgments

We thank Naim Rashid, Scott Van Buren, Yun Li, Jason Stein, Jeremy Simon, and Frank Konietschke for useful discussions.

