## Supplementary Tables and Figures for "Nonparametric expression analysis using inferential replicate counts"

### Supplementary Tables & Figures

| | DESeq2 | EBSeq | limma | swish | DESeq2 |
|---|---|---|---|---|---|
| | (counts w/ outlier) | | | | (original counts) |
| ENST00000488147.1 | 1.30E-05 | 5.31E-09 | 0.723 | 0.834 | 0.984 |
| ENST00000623083.4 | 3.49E-06 | 1.62E-09 | 0.758 | 0.847 | 0.727 |
| ENST00000599771.6 | 1.97E-04 | 1.46E-07 | 0.752 | 0.898 | 0.591 |

Supplementary Table 1: Outlier experiment for the bulk RNA-seq simulation with 6 samples per group, one batch. The table presents adjusted $p$ values for three null transcripts, in which each transcript had a single sample with an outlier count inserted, by multiplying the count by 1,000. The rightmost column gives the results of running *DESeq2* on the original counts, without the outlier inserted. Adjusted $p$ values less than 0.01 are underlined. *DESeq2* was run with its Cook's-distance-based outlier flagging turned off.

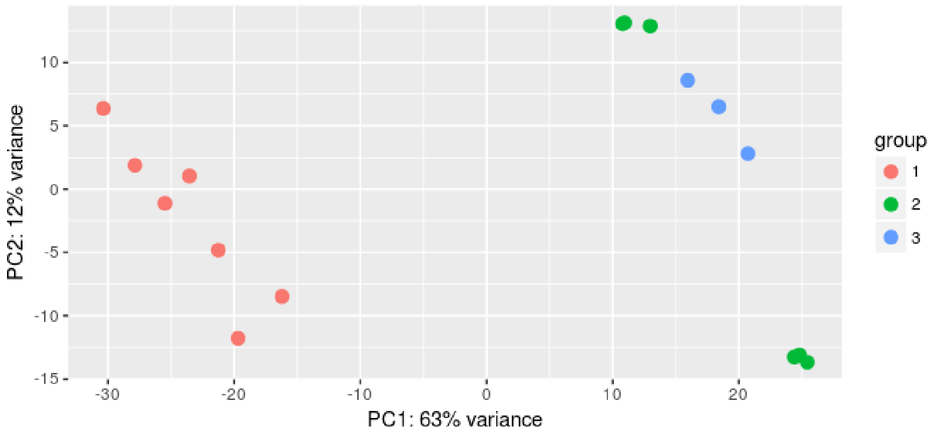


Supplementary Figure 1: PCA plot of variance stabilized counts from the *Arabidopsis* highly replicated dataset, for the WT condition. Batch 1 (red points) was separated from batches 2 and 3 (green and blue points), and so partitions of the data were chosen such that the number of samples from batch 1 would be balanced across the random splits.

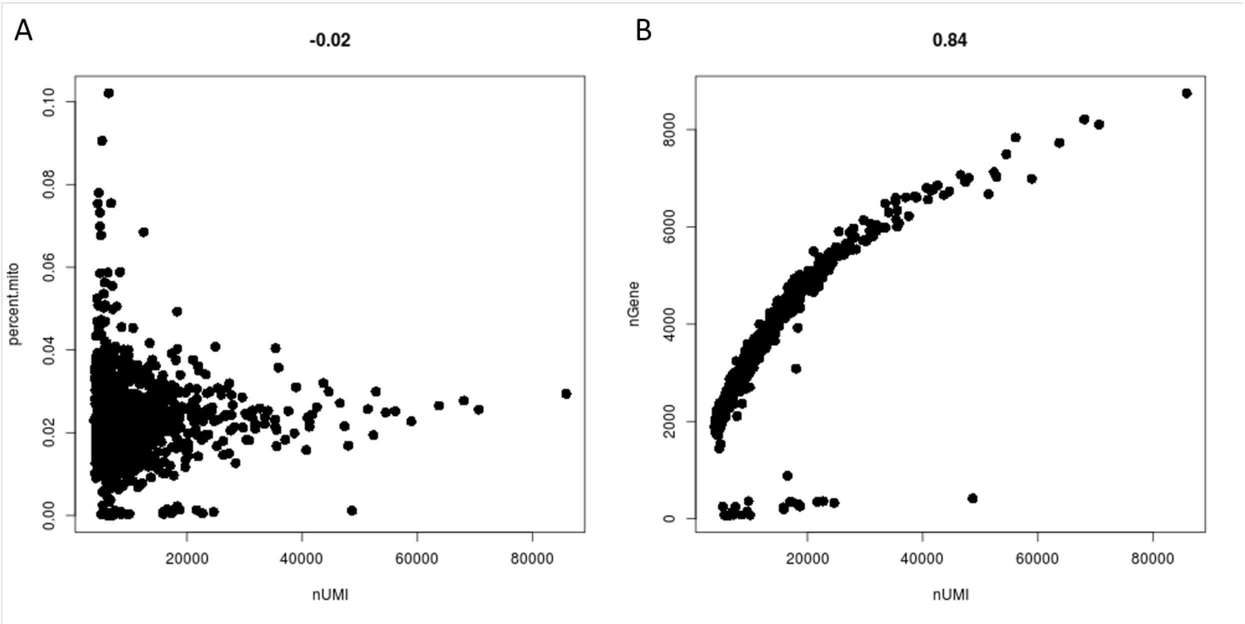


Supplementary Figure 2: *Seurat* quality control scatterplots for cells in the developing mouse brain scRNA-seq dataset. (A) Percentage of mitochondrial genes detected over molecule counts and (B) detected gene counts over molecule counts. The plots were used to detect cells of low quality with either high percentage of mitochondrial genes detected or potential multiplets with outlying number of genes detected given the total molecular count.

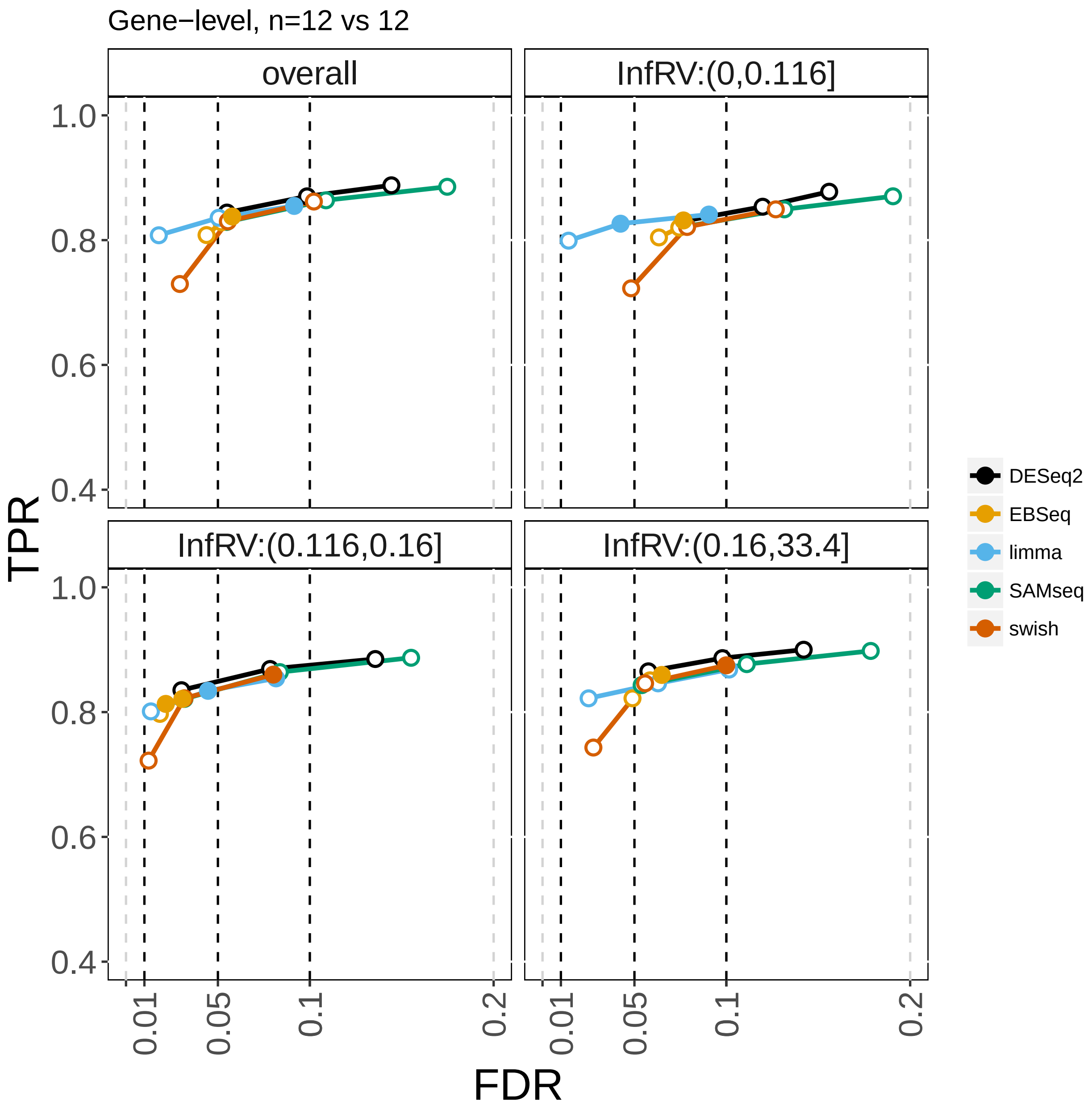


Supplementary Figure 3: True positive rate (y-axis) over false discovery rate (x-axis) for DGE analysis with two batches of samples, with 12 samples in each of the two conditions (bulk RNA-seq simulated dataset). The first panel depicts overall performance, while the subsequent three panels depict the genes stratified into thirds by InfRV averaged over samples. *sleuth* is included in the following figure.

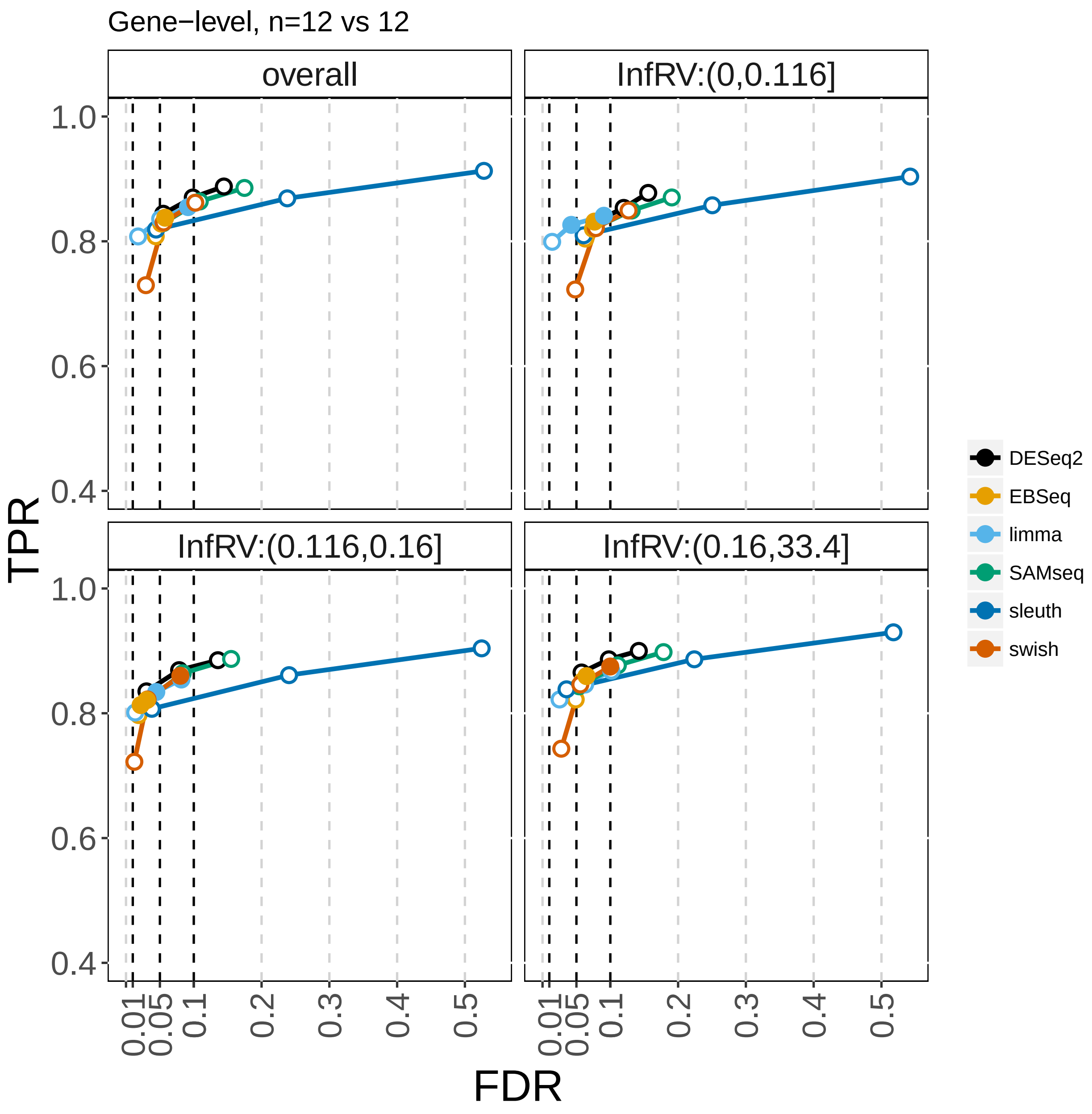


Supplementary Figure 4: True positive rate (y-axis) over false discovery rate (x-axis) for DGE analysis with two batches of samples, with 12 samples in each of the two conditions (bulk RNA-seq simulated dataset).

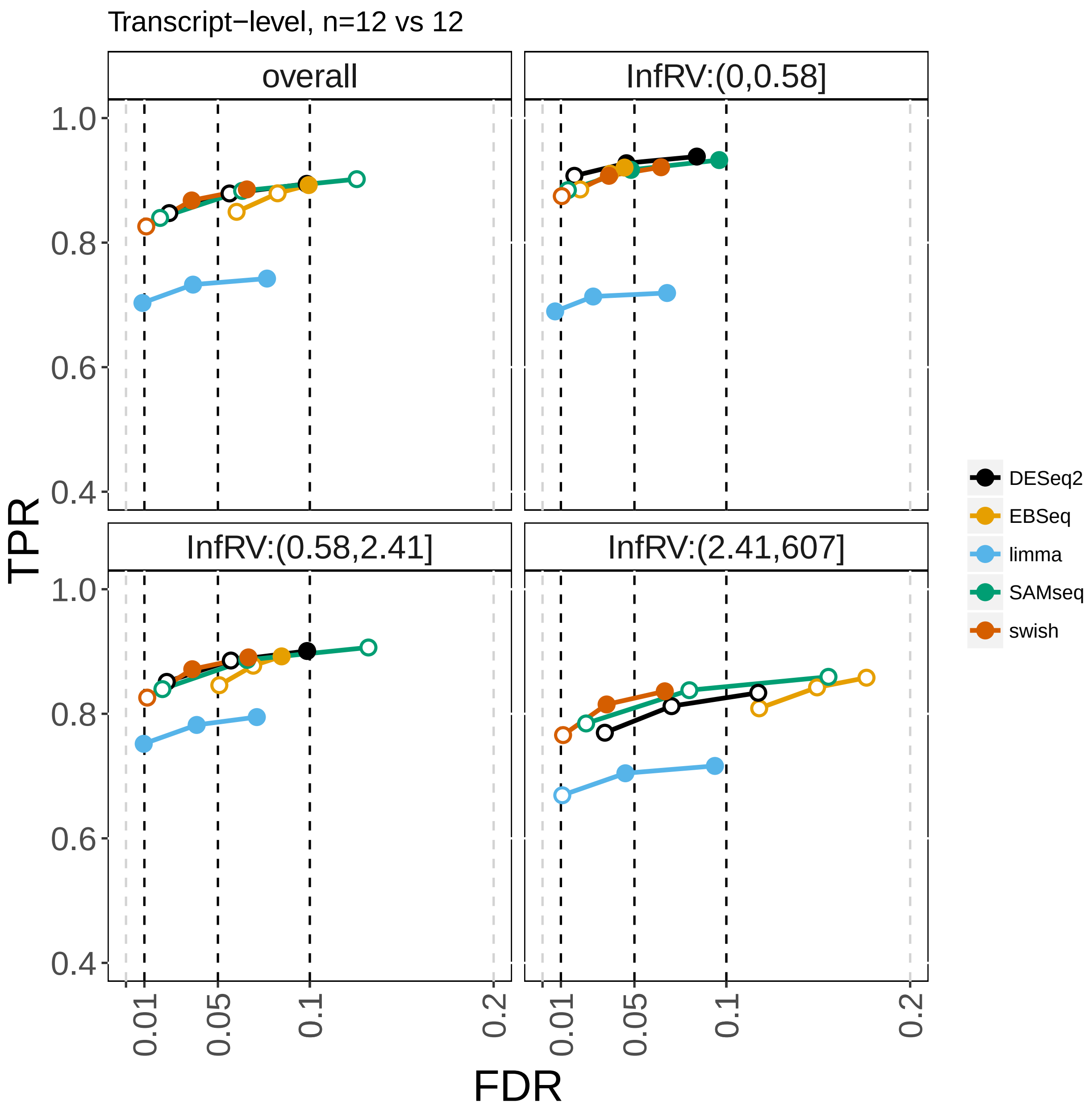


Supplementary Figure 5: True positive rate (y-axis) over false discovery rate (x-axis) for DTE analysis with two batches of samples, with 12 samples in each of the two conditions (bulk RNA-seq simulated dataset). *sleuth* is included in the following figure.

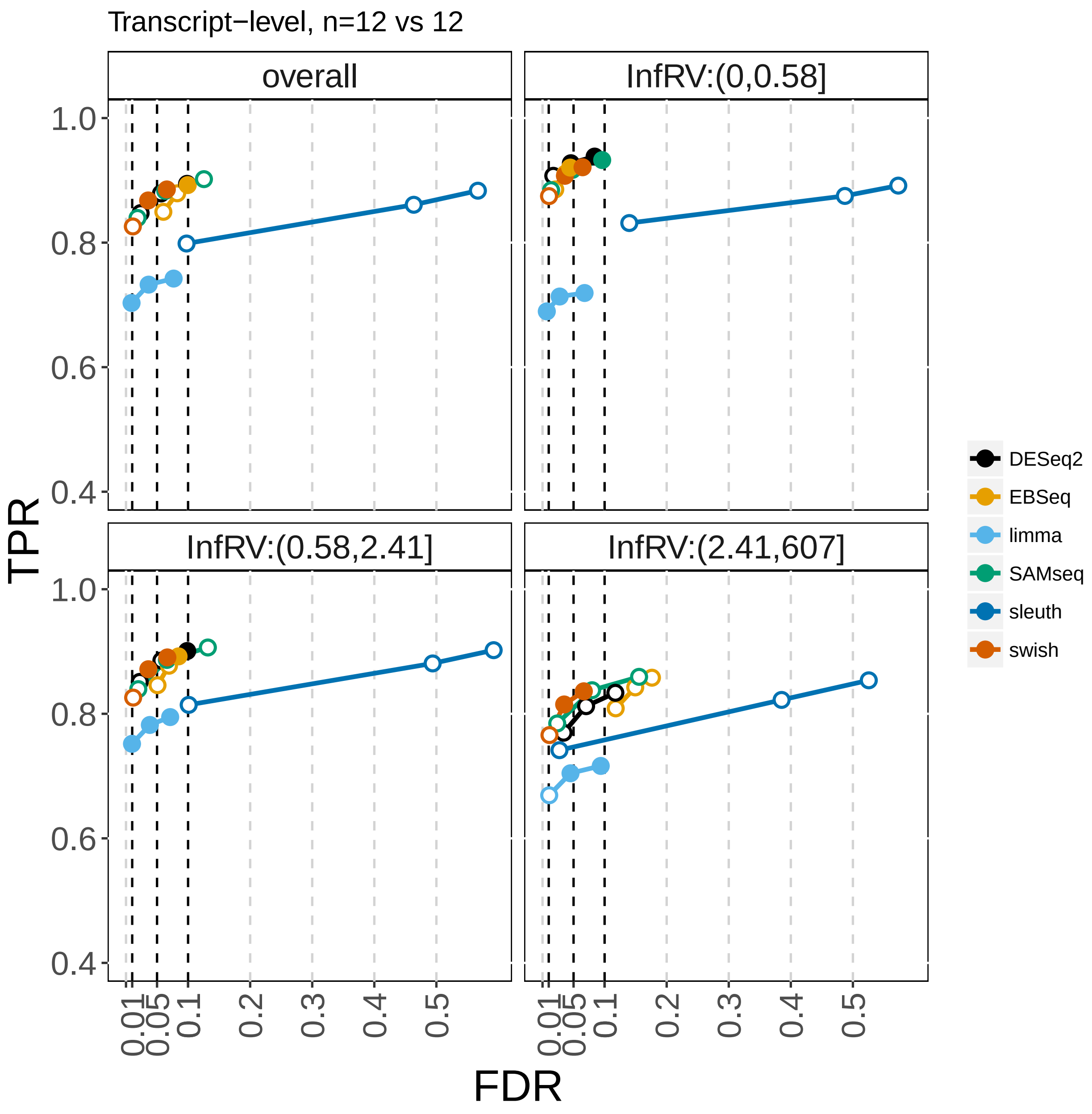


Supplementary Figure 6: True positive rate (y-axis) over false discovery rate (x-axis) for DTE analysis with two batches of samples, with 12 samples in each of the two conditions (bulk RNA-seq simulated dataset).

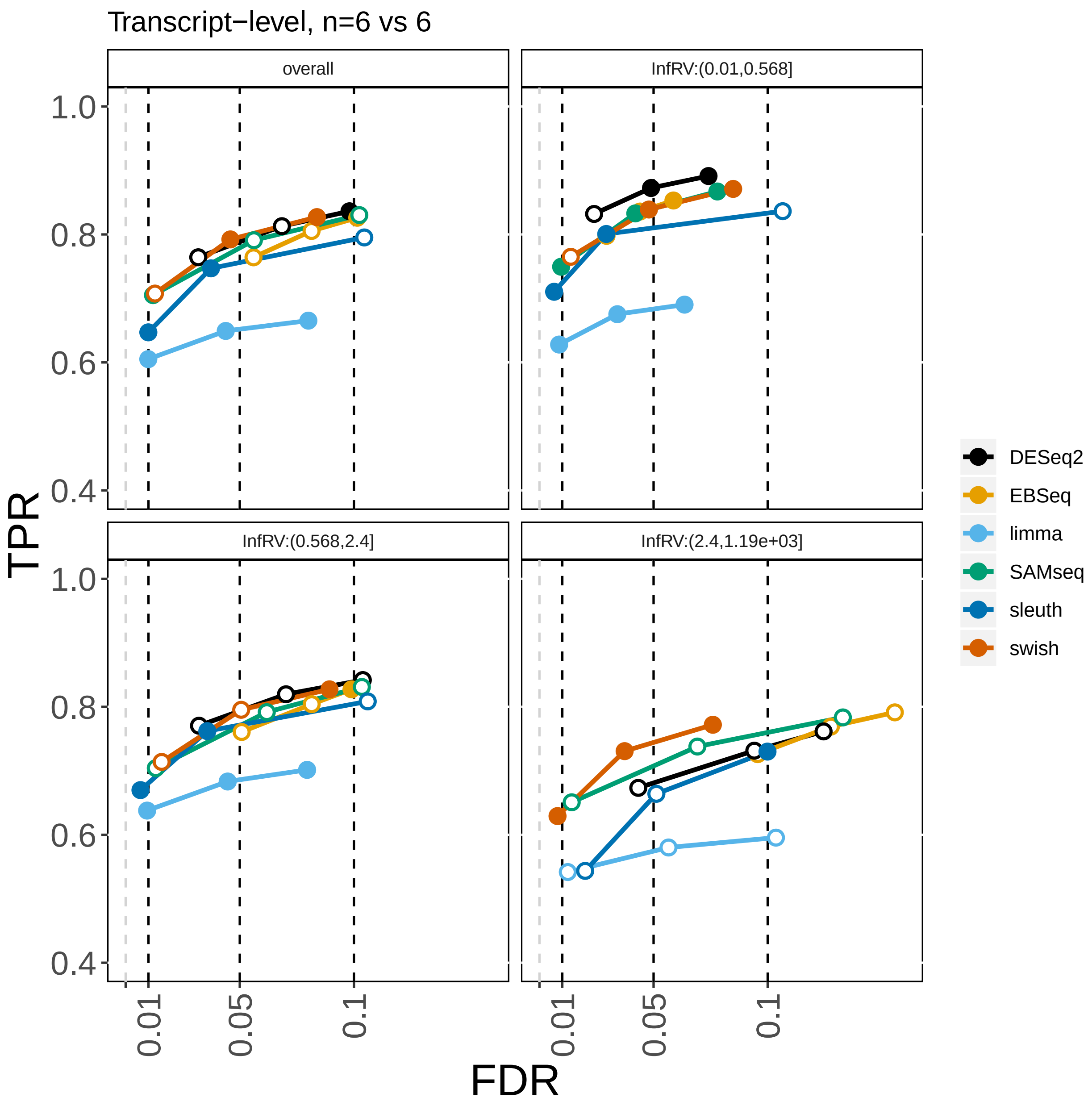


Supplementary Figure 7: Effect of filtering on sensitivity of *limma*. True positive rate (y-axis) over false discovery rate (x-axis) for DTE analysis with one batch of samples, with 6 samples in each of the two conditions (bulk RNA-seq simulated dataset.) In this analysis, *limma* was run with a minimum count of 3 for at least 6 samples, where the default is a minimum count of 10, and with a minimum total count of 5 across all samples, where the default is 15. The sensitivity is higher compared to Figure 3.

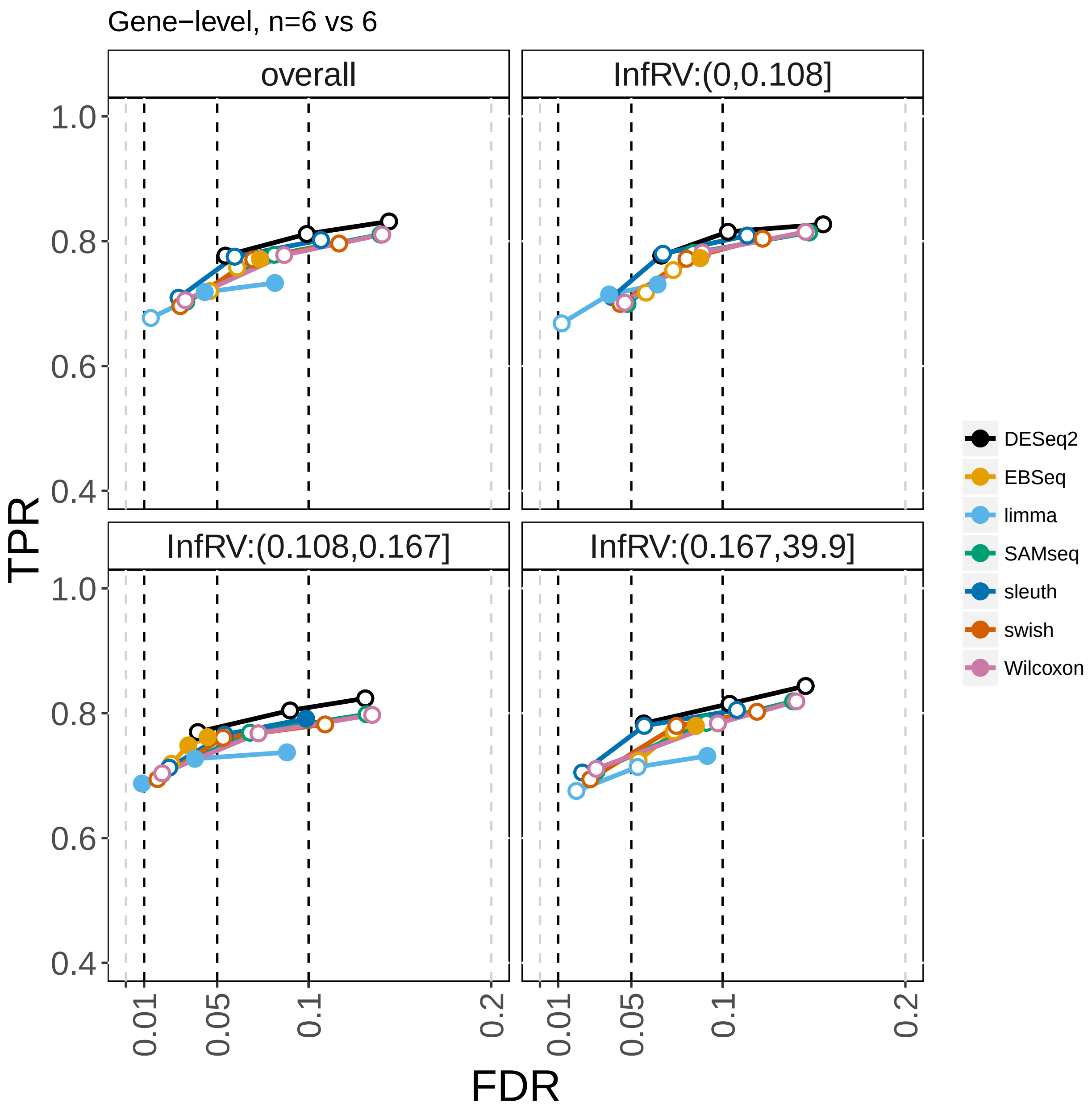


Supplementary Figure 8: True positive rate (y-axis) over false discovery rate (x-axis) for DGE analysis with one batch of samples, including the Wilcoxon test.

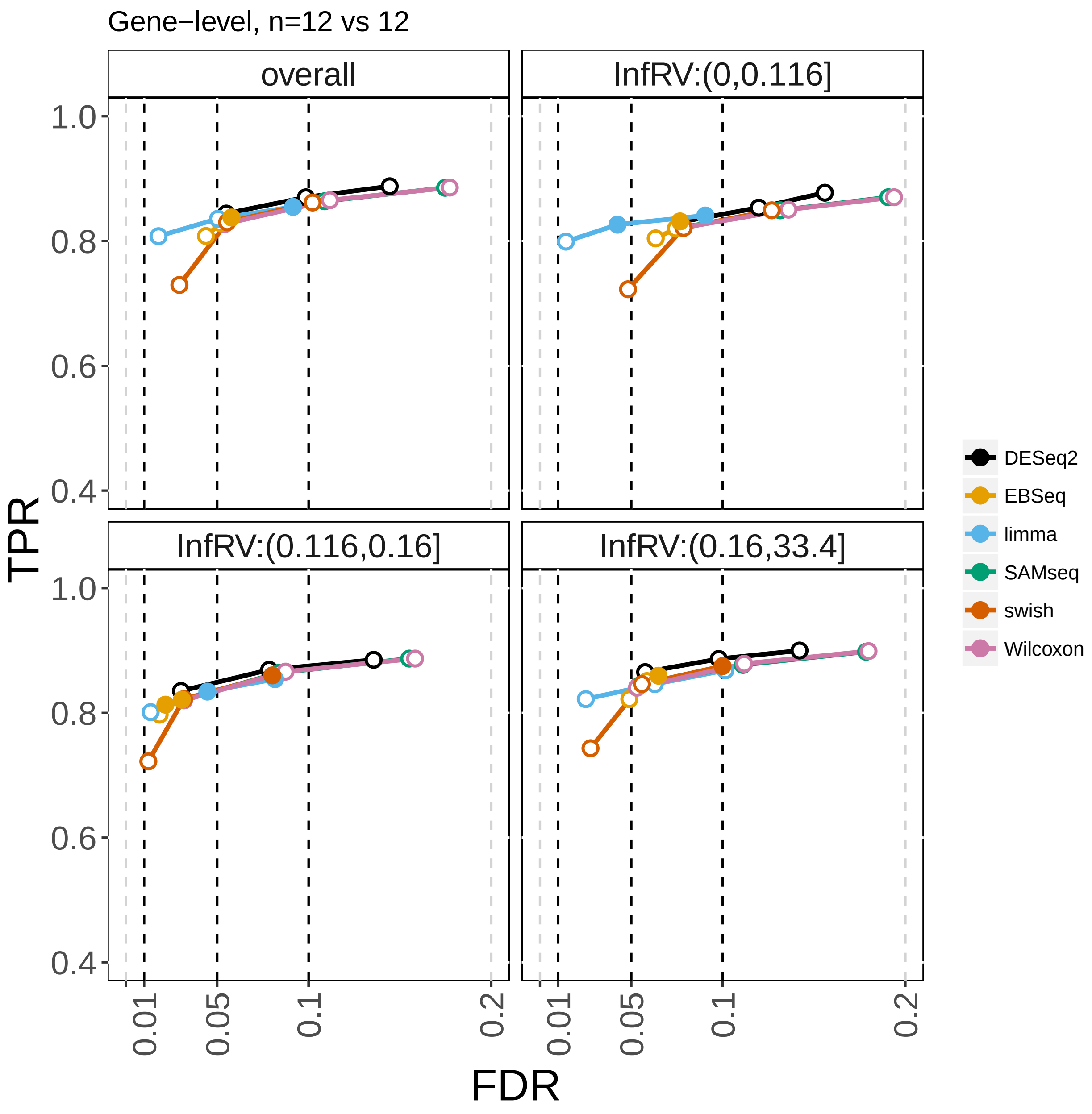


Supplementary Figure 9: True positive rate (y-axis) over false discovery rate (x-axis) for DGE analysis with two batches of samples, including the Wilcoxon test.

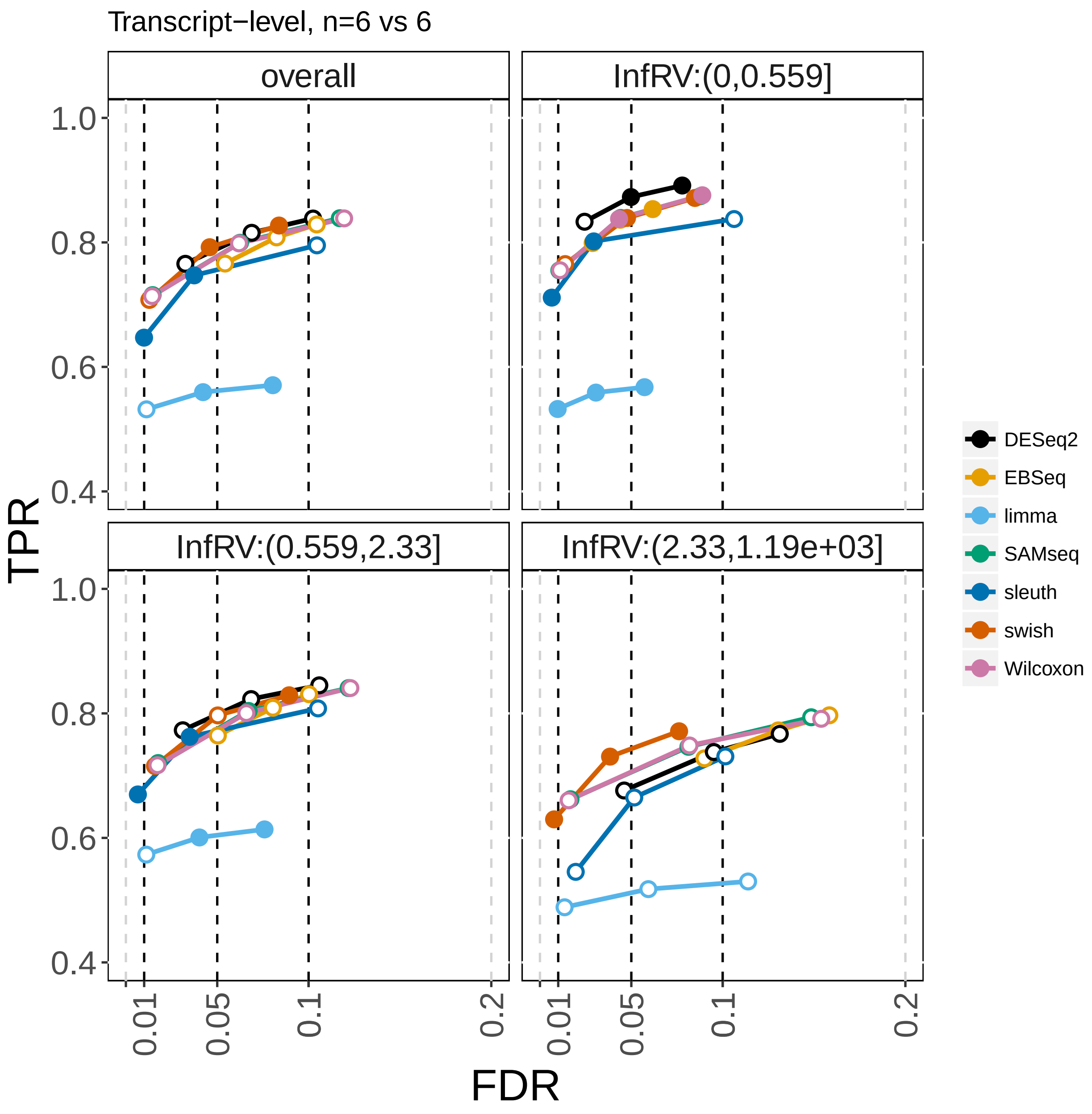


Supplementary Figure 10: True positive rate (y-axis) over false discovery rate (x-axis) for DGE analysis with one batch of samples, including the Wilcoxon test.

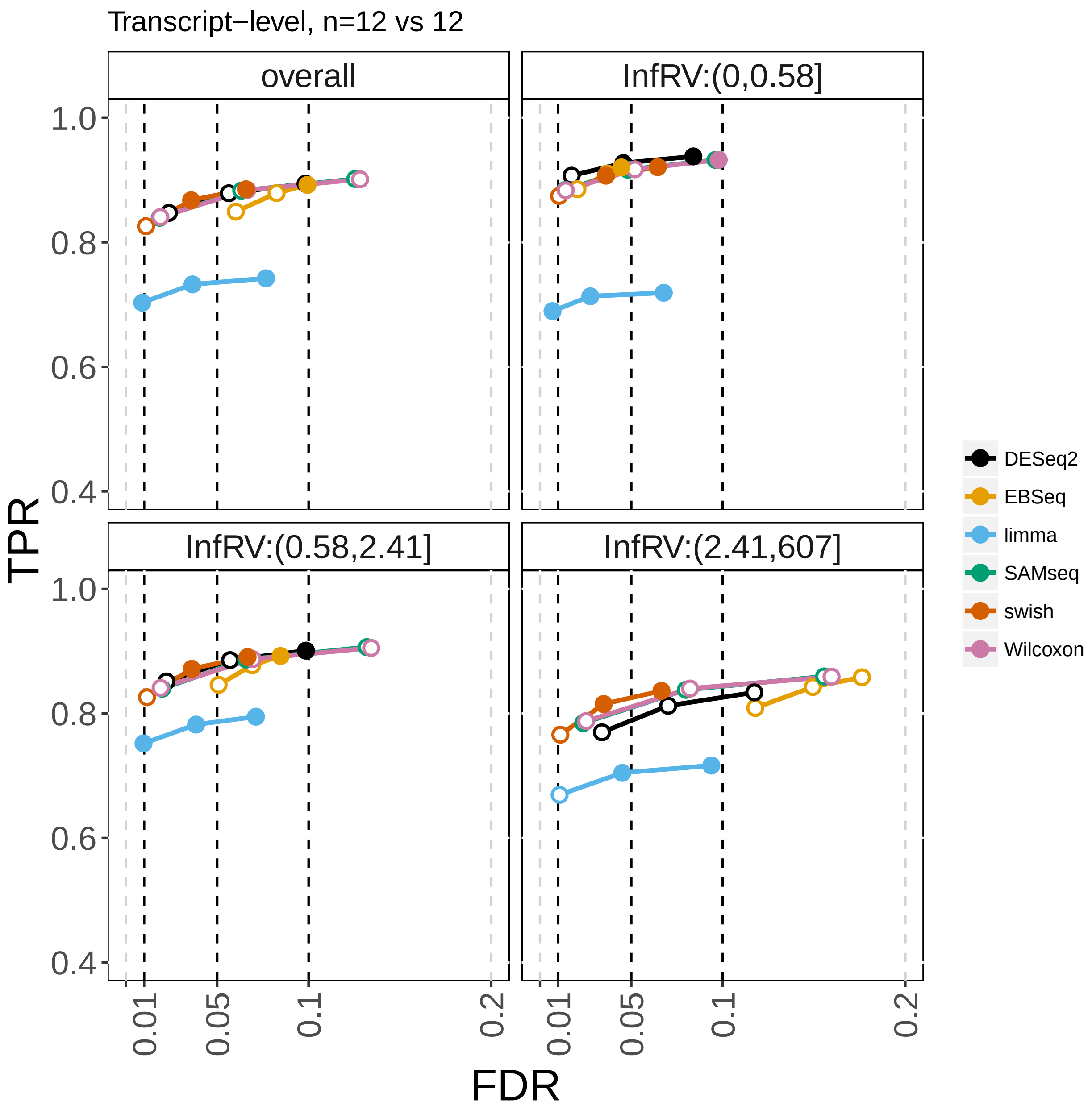


Supplementary Figure 11: True positive rate (y-axis) over false discovery rate (x-axis) for DGE analysis with two batches of samples, including the Wilcoxon test.

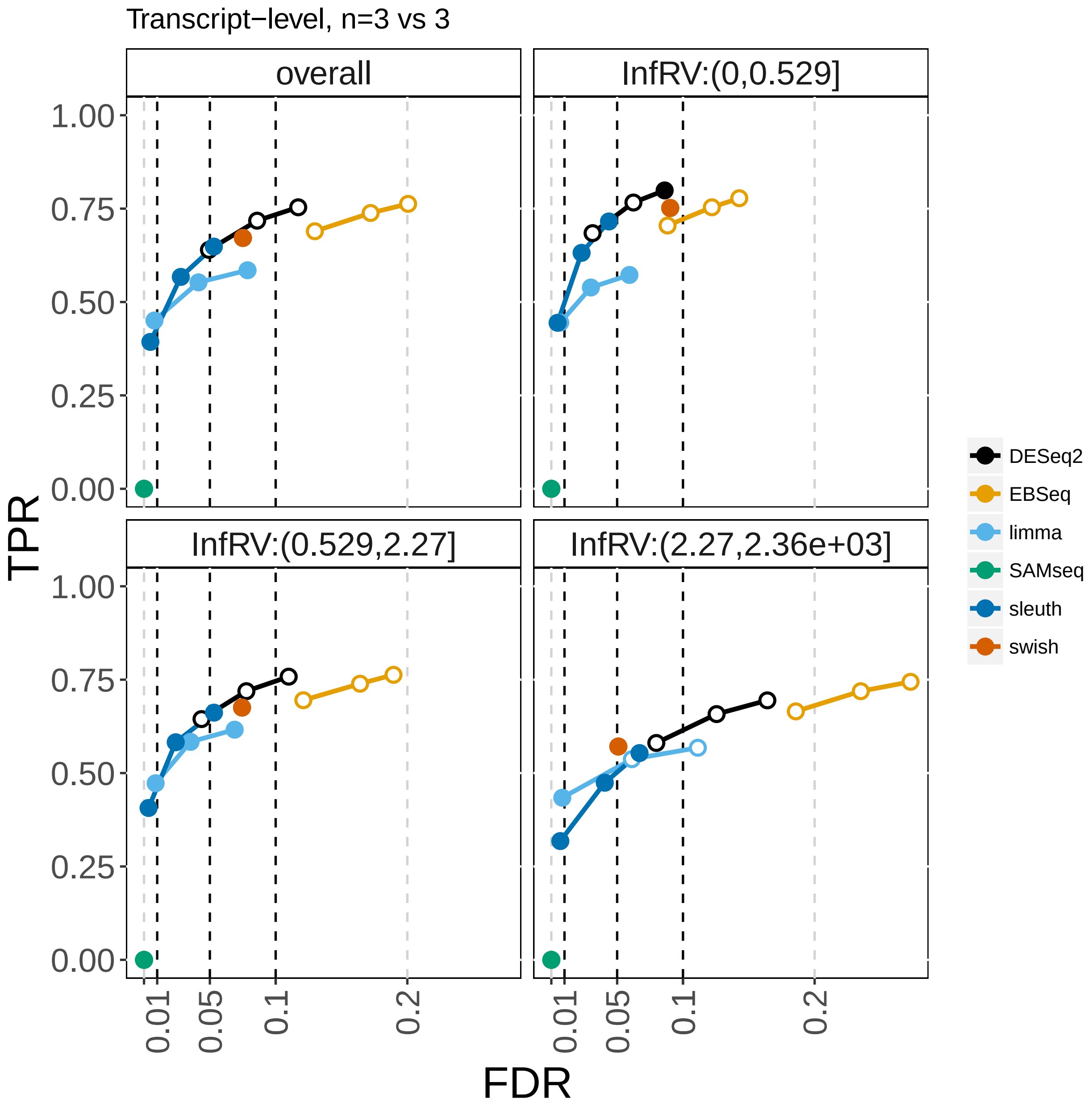


Supplementary Figure 12: True positive rate (y-axis) over false discovery rate (x-axis) for DTE analysis with one batch of samples.

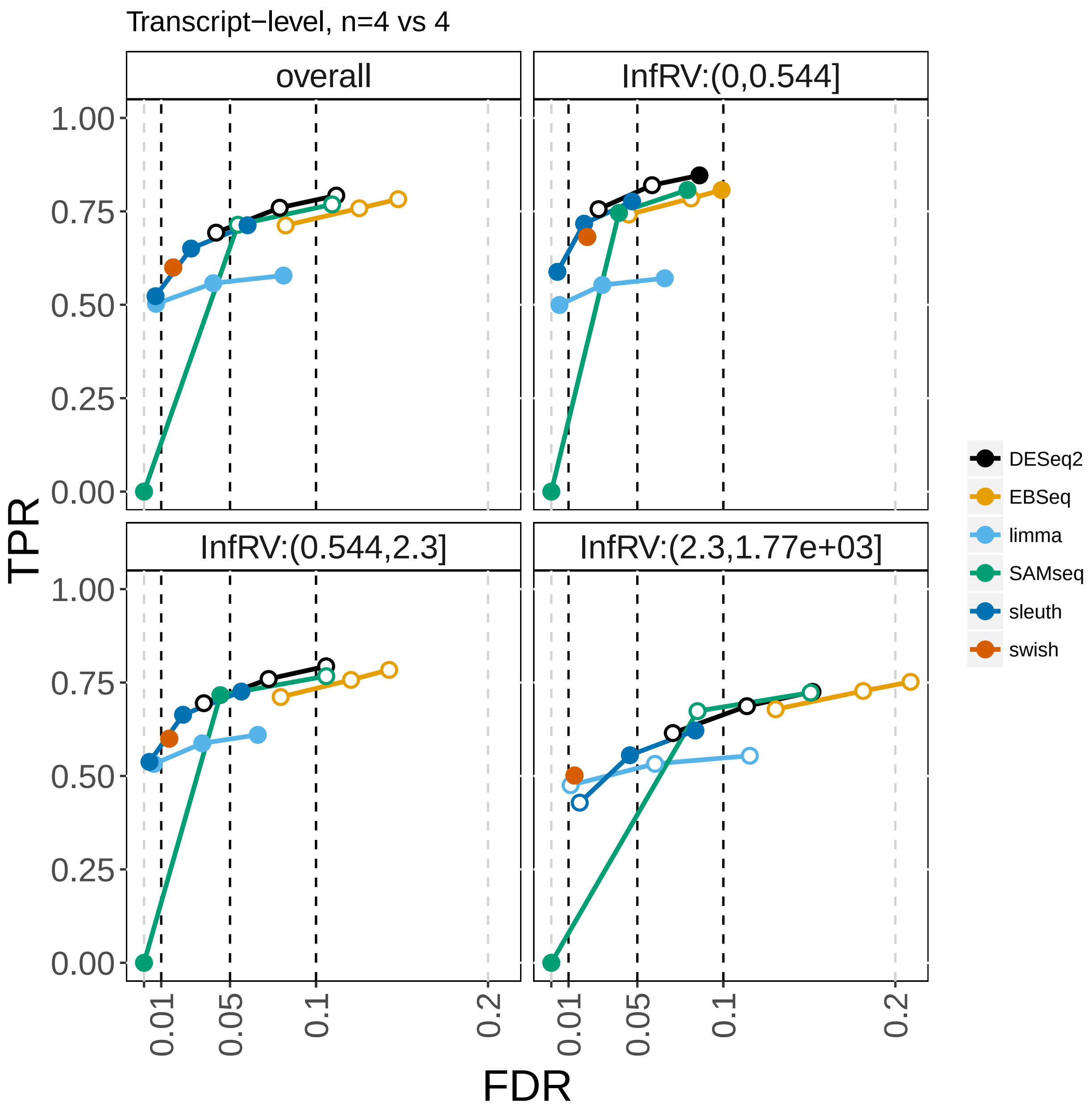


Supplementary Figure 13: True positive rate (y-axis) over false discovery rate (x-axis) for DTE analysis with one batch of samples.

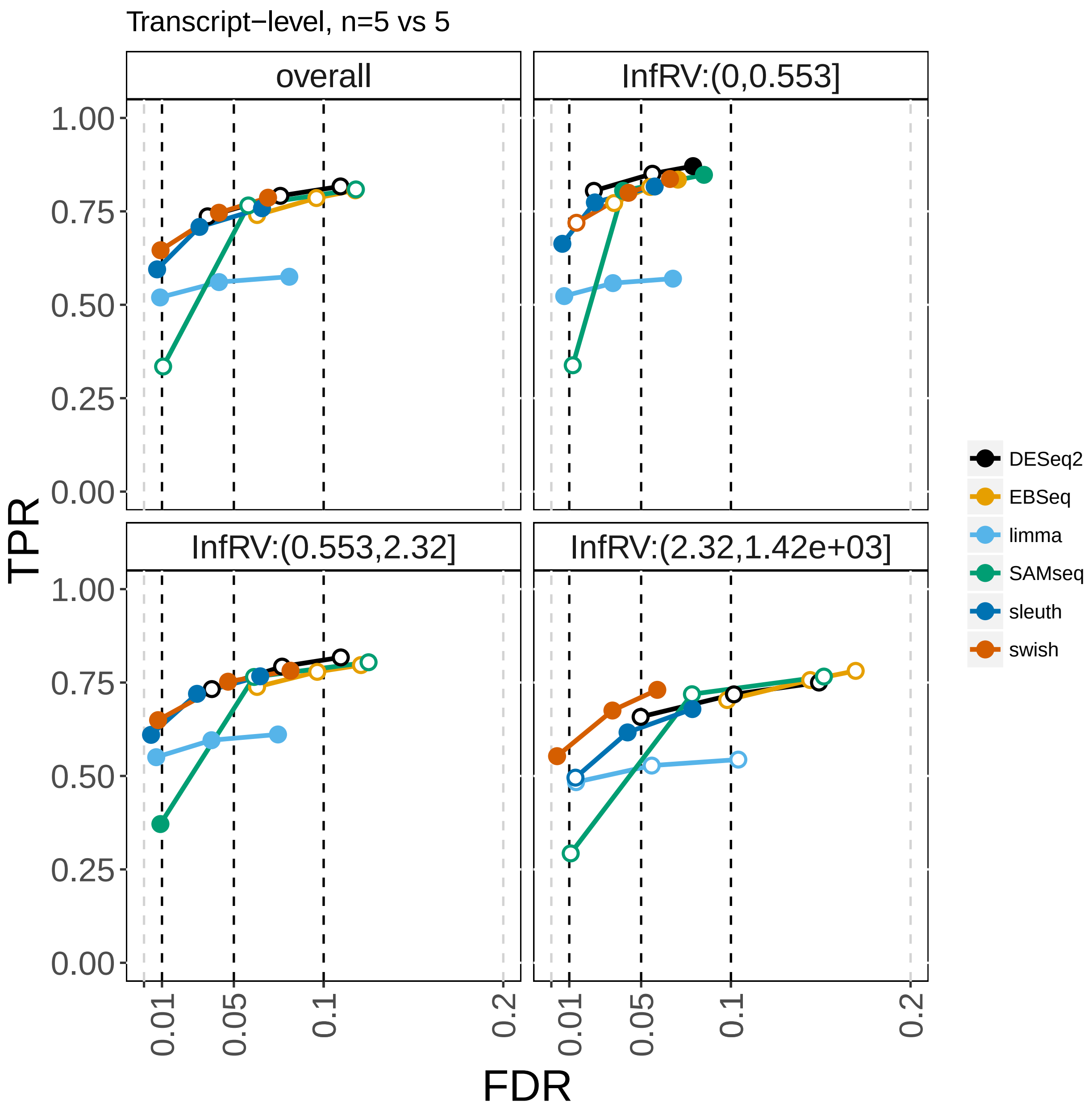


Supplementary Figure 14: True positive rate (y-axis) over false discovery rate (x-axis) for DTE analysis with one batch of samples.

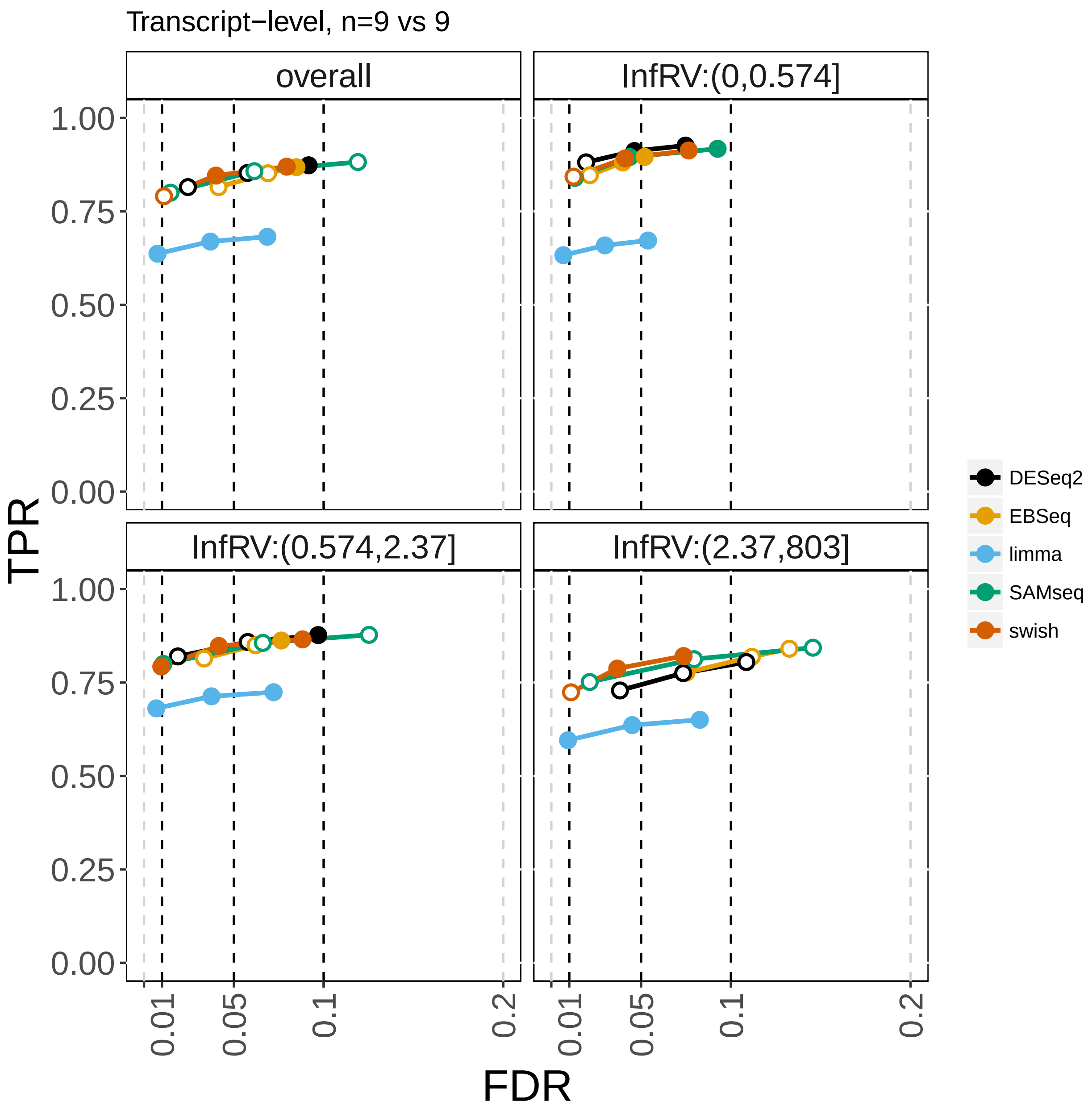


Supplementary Figure 15: True positive rate (y-axis) over false discovery rate (x-axis) for DTE analysis with one batch of samples.

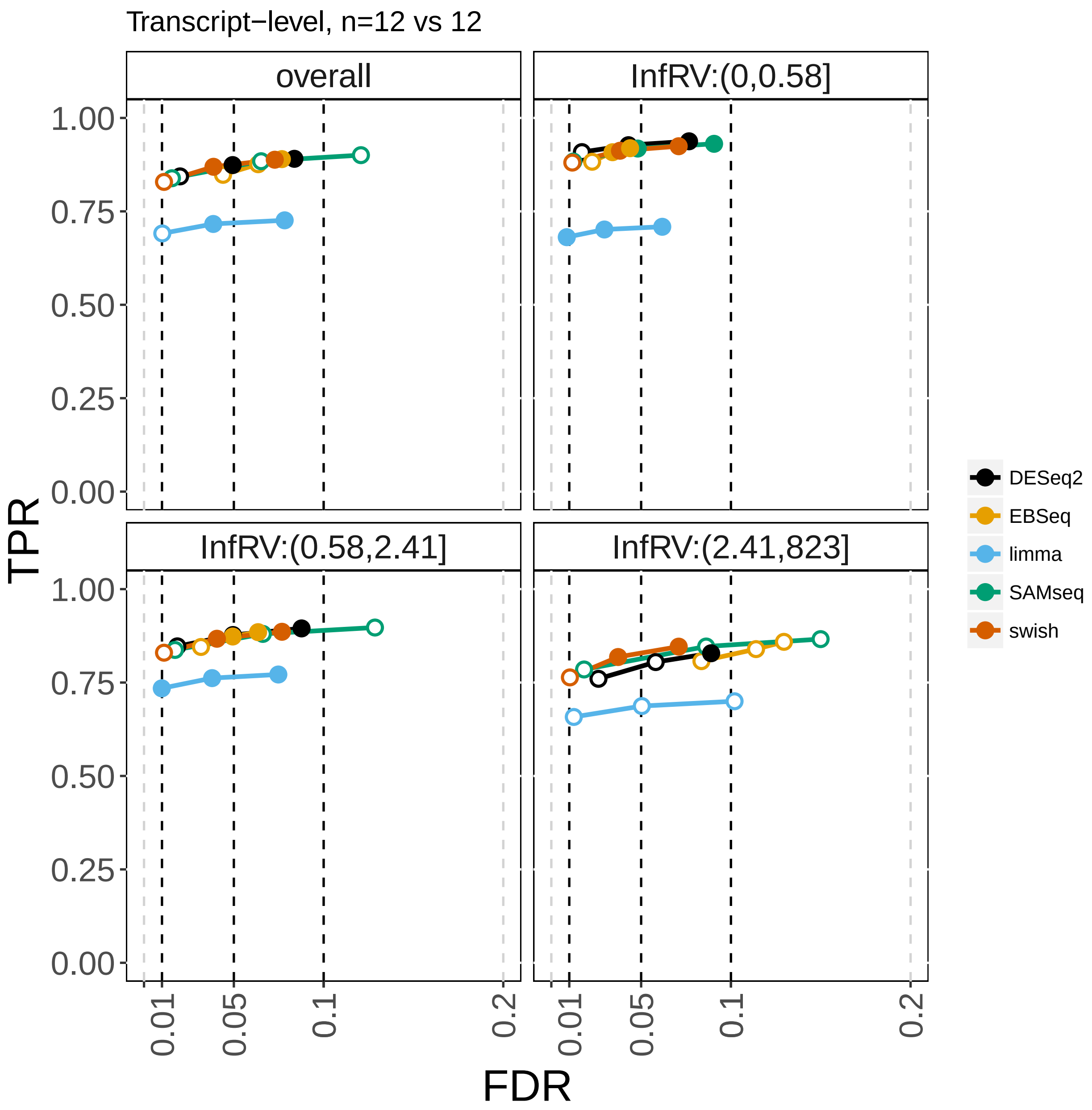


Supplementary Figure 16: True positive rate (y-axis) over false discovery rate (x-axis) for DTE analysis with one batch of samples.

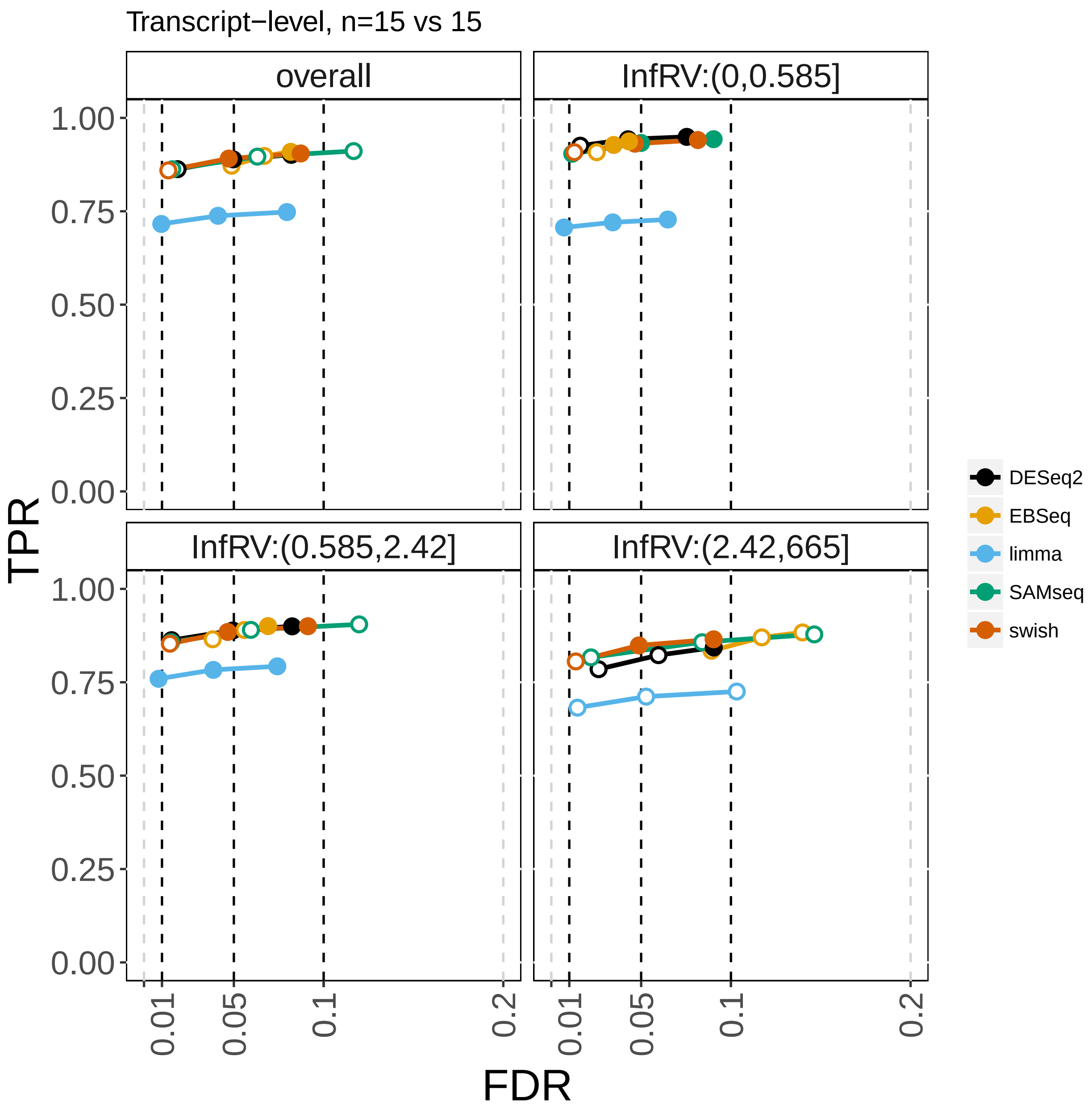


Supplementary Figure 17: True positive rate (y-axis) over false discovery rate (x-axis) for DTE analysis with one batch of samples.

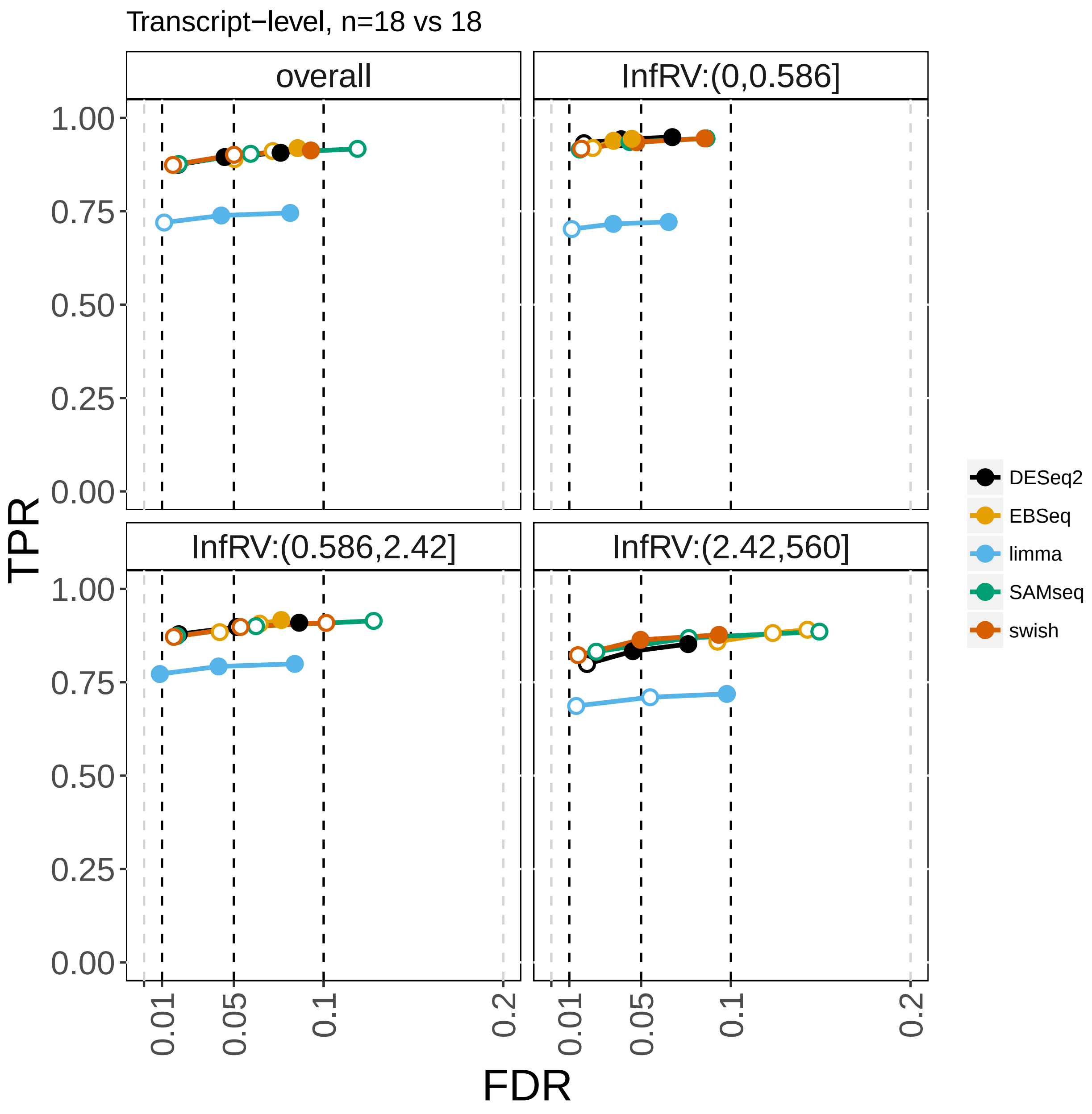


Supplementary Figure 18: True positive rate (y-axis) over false discovery rate (x-axis) for DTE analysis with one batch of samples.

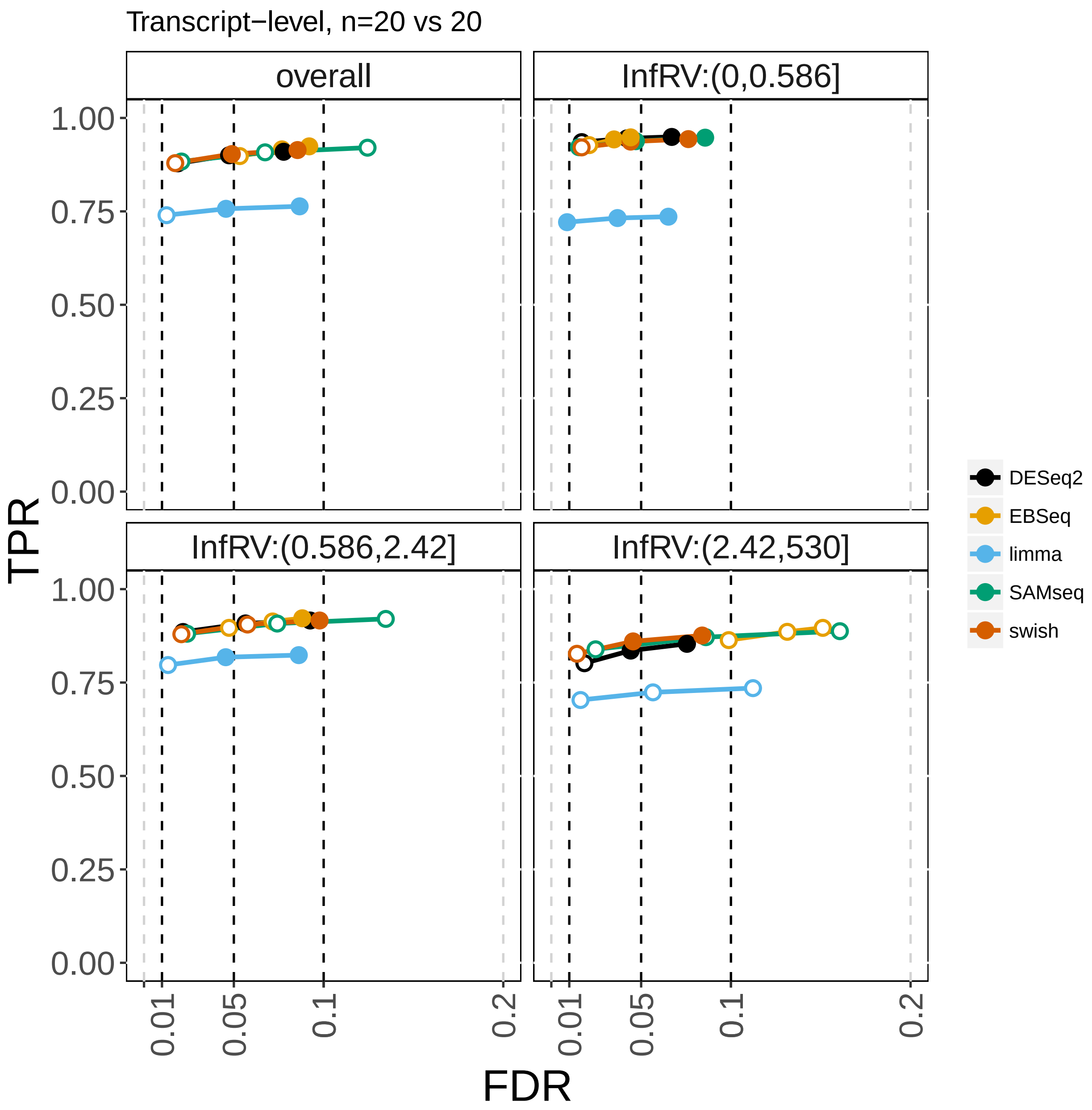


Supplementary Figure 19: True positive rate (y-axis) over false discovery rate (x-axis) for DTE analysis with one batch of samples.

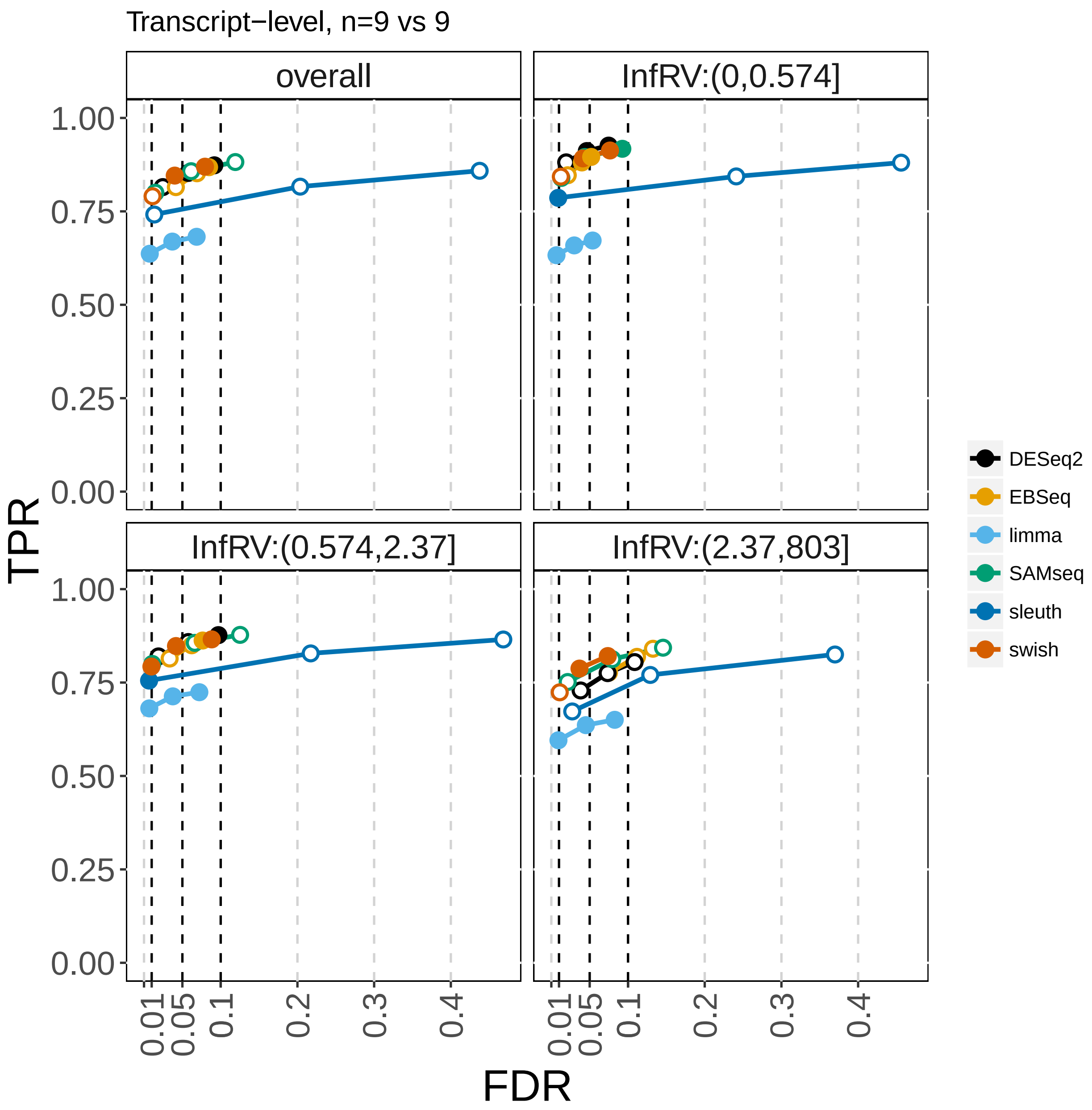


Supplementary Figure 20: True positive rate (y-axis) over false discovery rate (x-axis) for DTE analysis with one batch of samples, including *sleuth*.

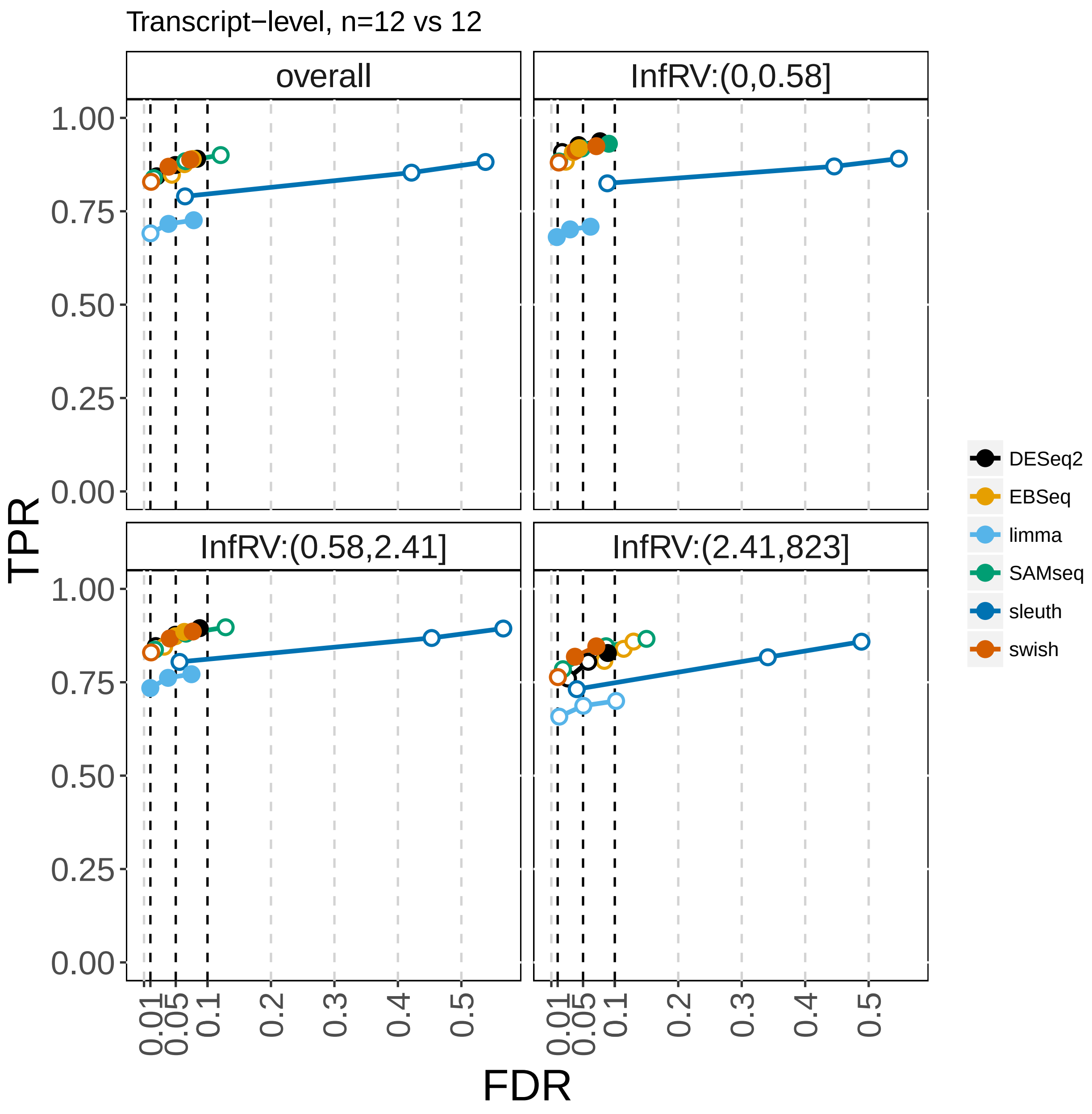


Supplementary Figure 21: True positive rate (y-axis) over false discovery rate (x-axis) for DTE analysis with one batch of samples, including *sleuth*.

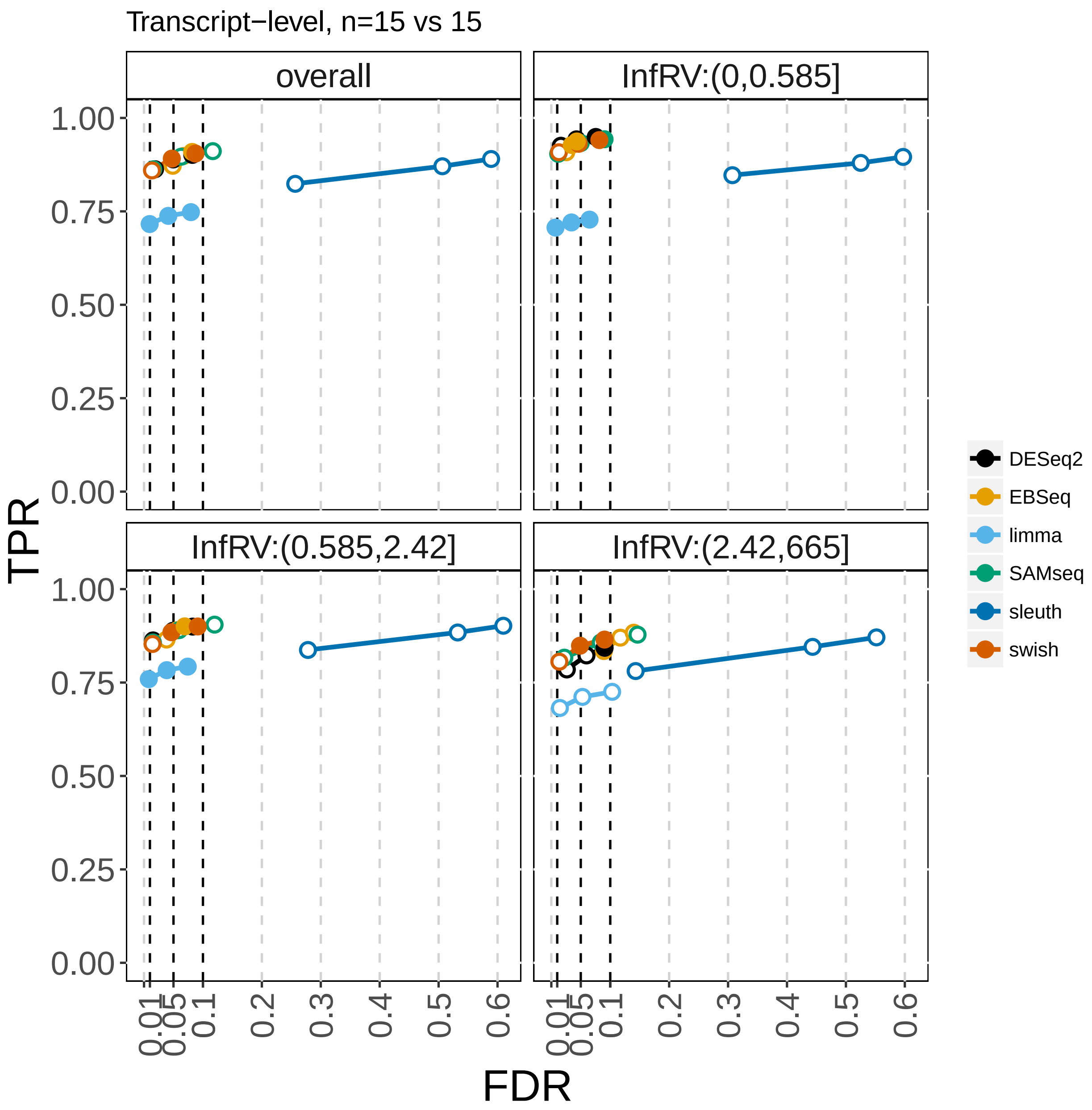


Supplementary Figure 22: True positive rate (y-axis) over false discovery rate (x-axis) for DTE analysis with one batch of samples, including *sleuth*.

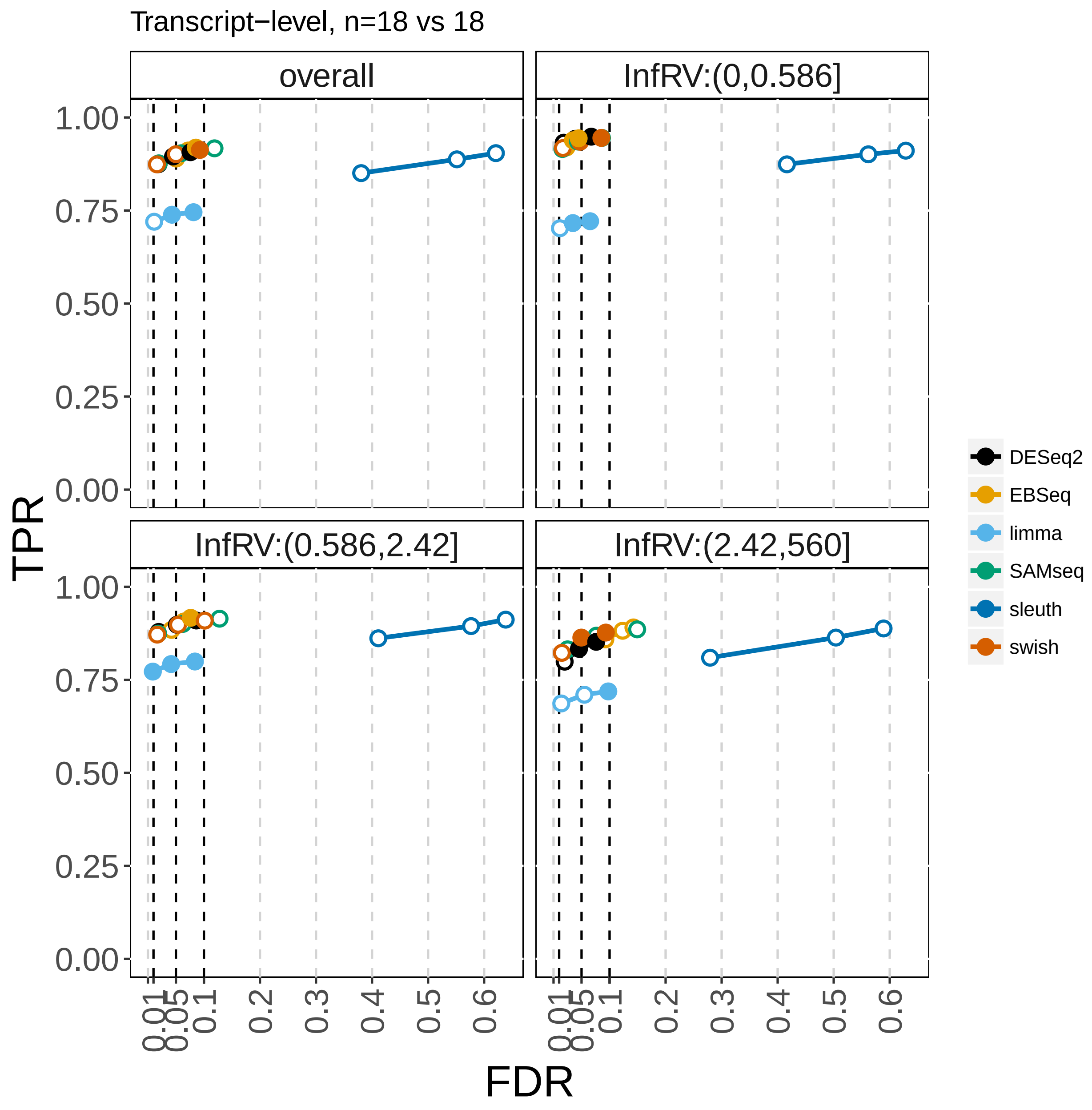


Supplementary Figure 23: True positive rate (y-axis) over false discovery rate (x-axis) for DTE analysis with one batch of samples, including *sleuth*.

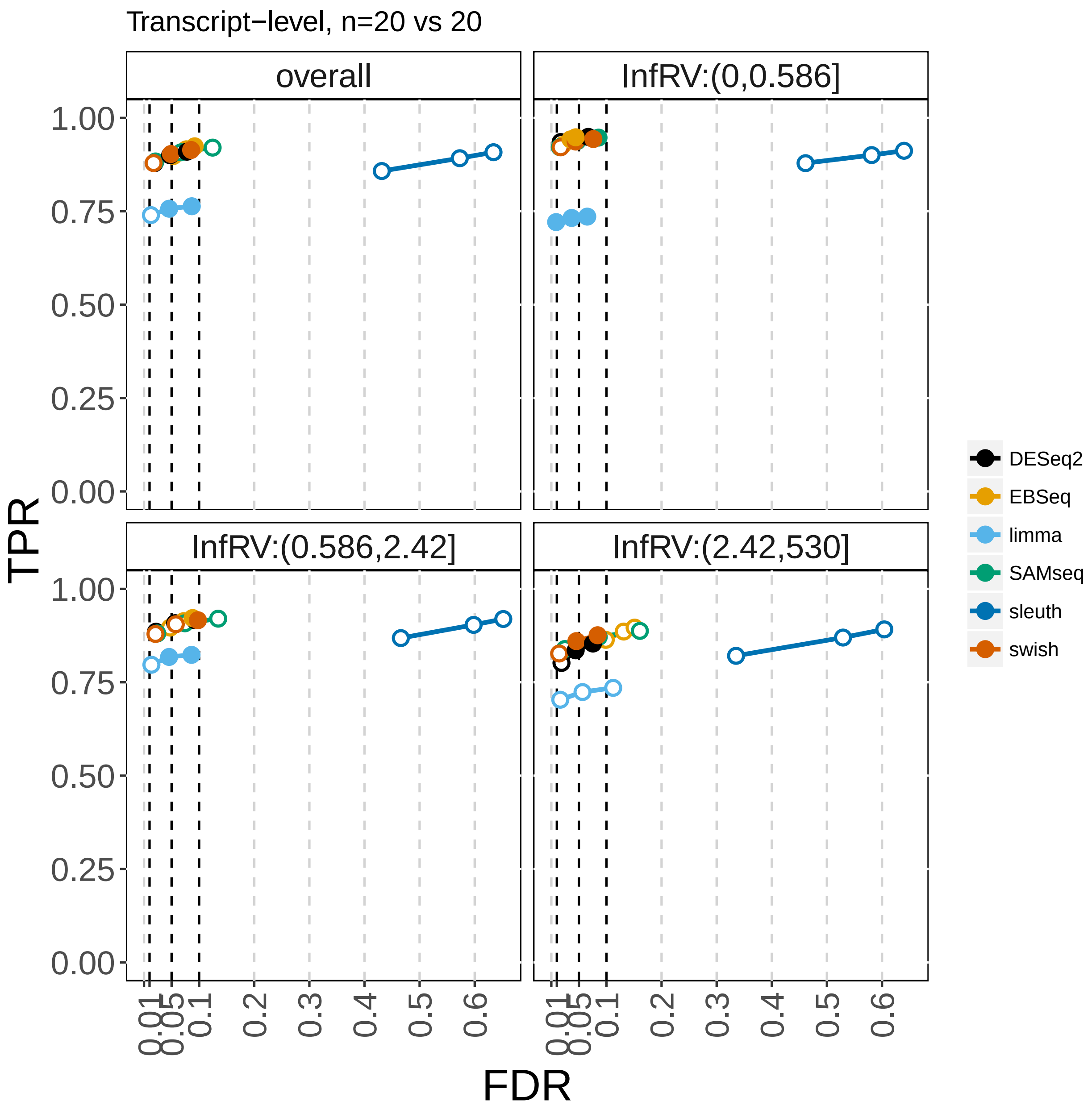


Supplementary Figure 24: True positive rate (y-axis) over false discovery rate (x-axis) for DTE analysis with one batch of samples, including *sleuth*.

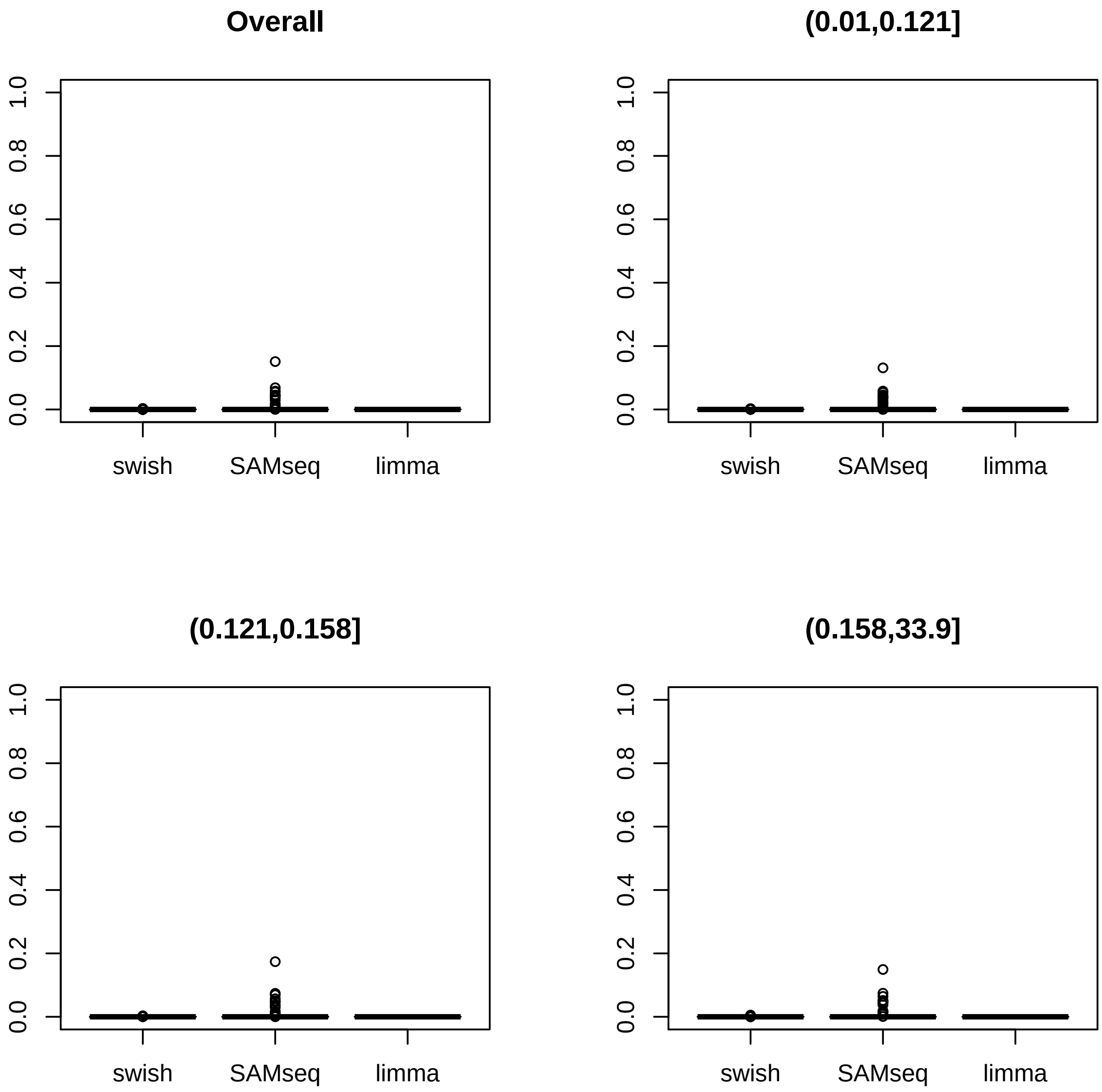


Supplementary Figure 25: False positive rate of *swish*, *SAMseq*, and *limma* over 100 repititions for yeast data at sample size 5. Panels provide overall FPR and FPR for features divided into thirds by InfRV.

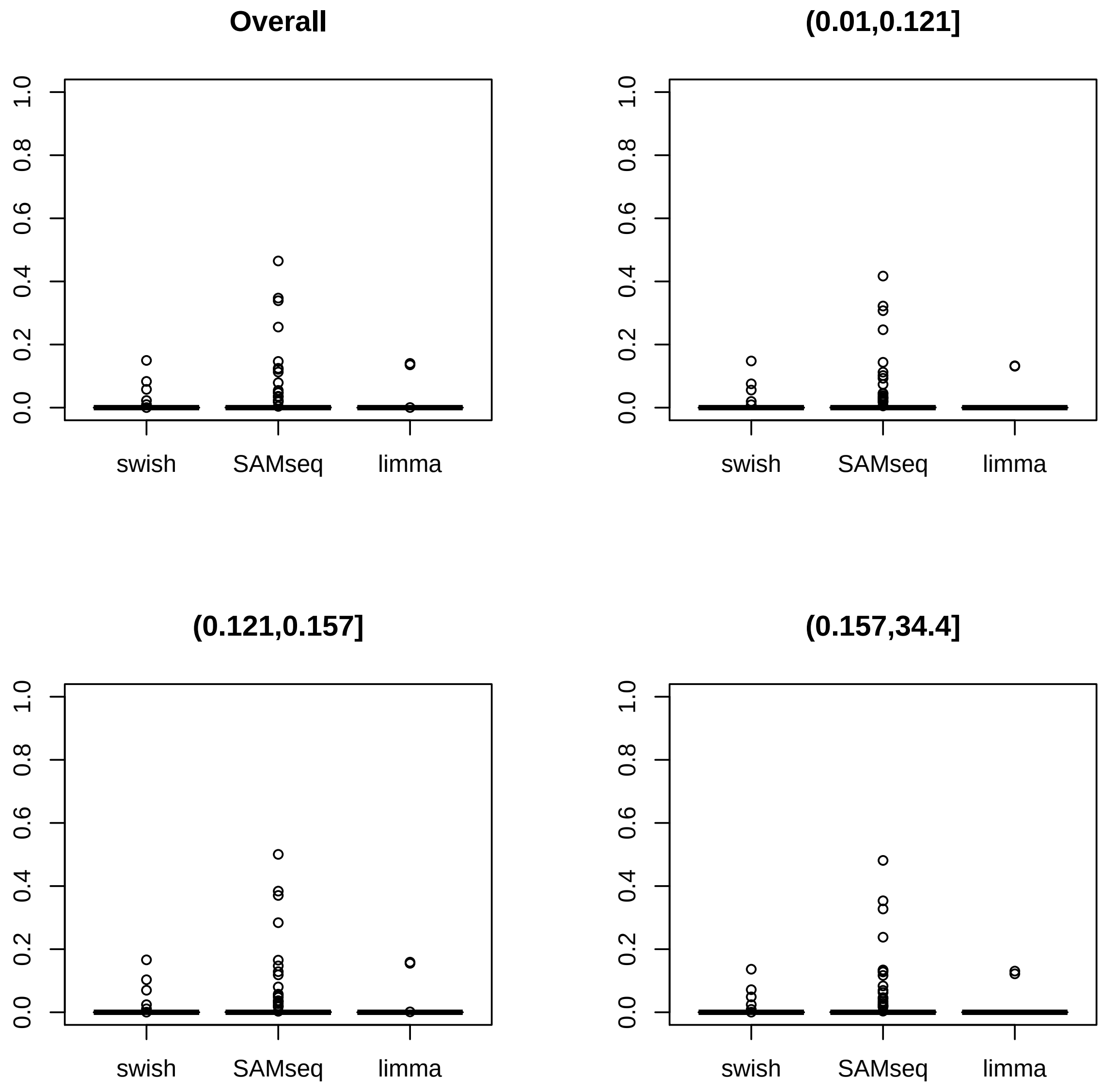


Supplementary Figure 26: False positive rate of *swish*, *SAMseq*, and *limma* over 100 repititions for yeast data at sample size 10. Panels provide overall FPR and FPR for features divided into thirds by InfRV.

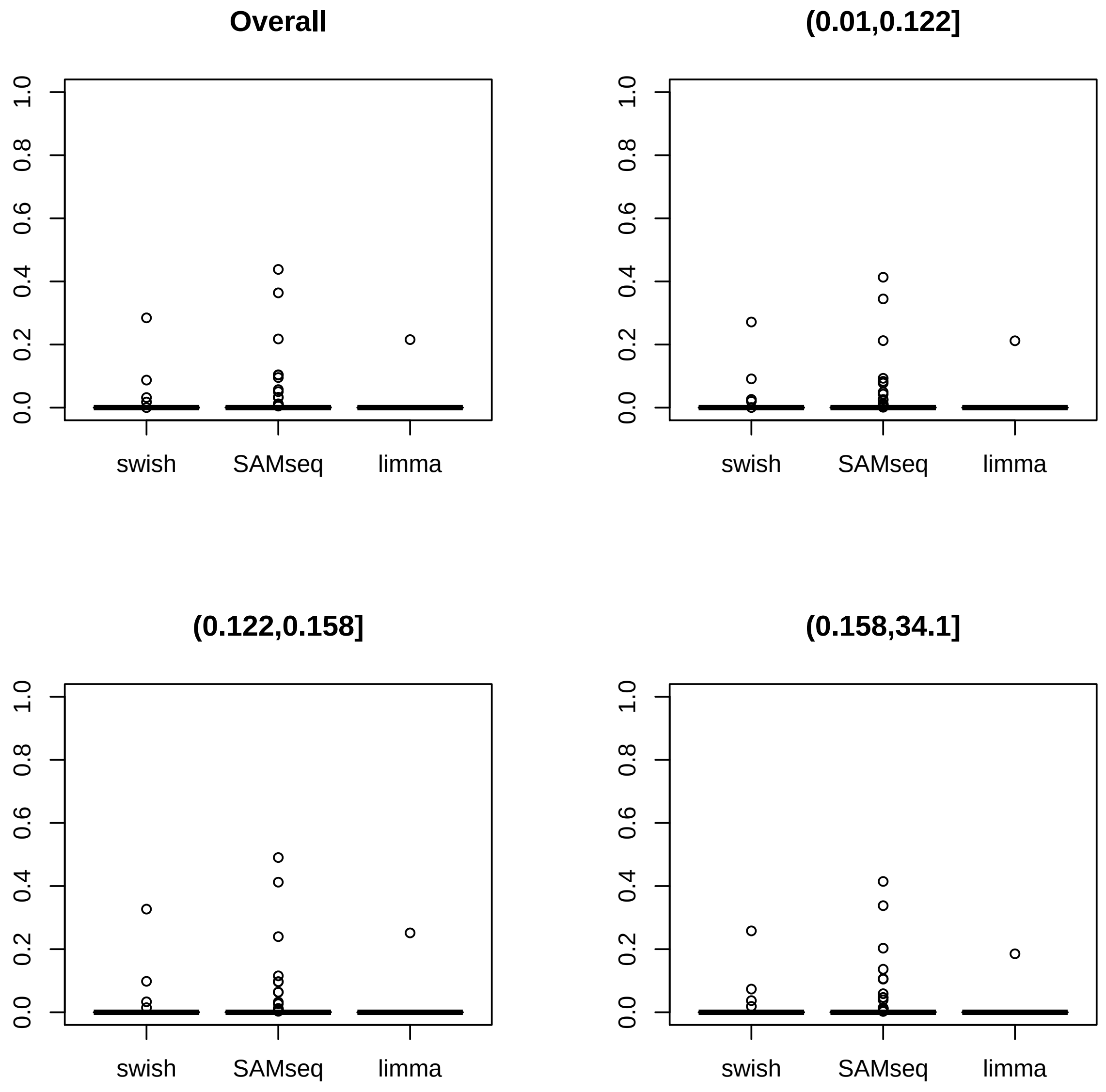


Supplementary Figure 27: False positive rate of *swish*, *SAMseq*, and *limma* over 100 repititions for yeast data at sample size 15. Panels provide overall FPR and FPR for features divided into thirds by InfRV.

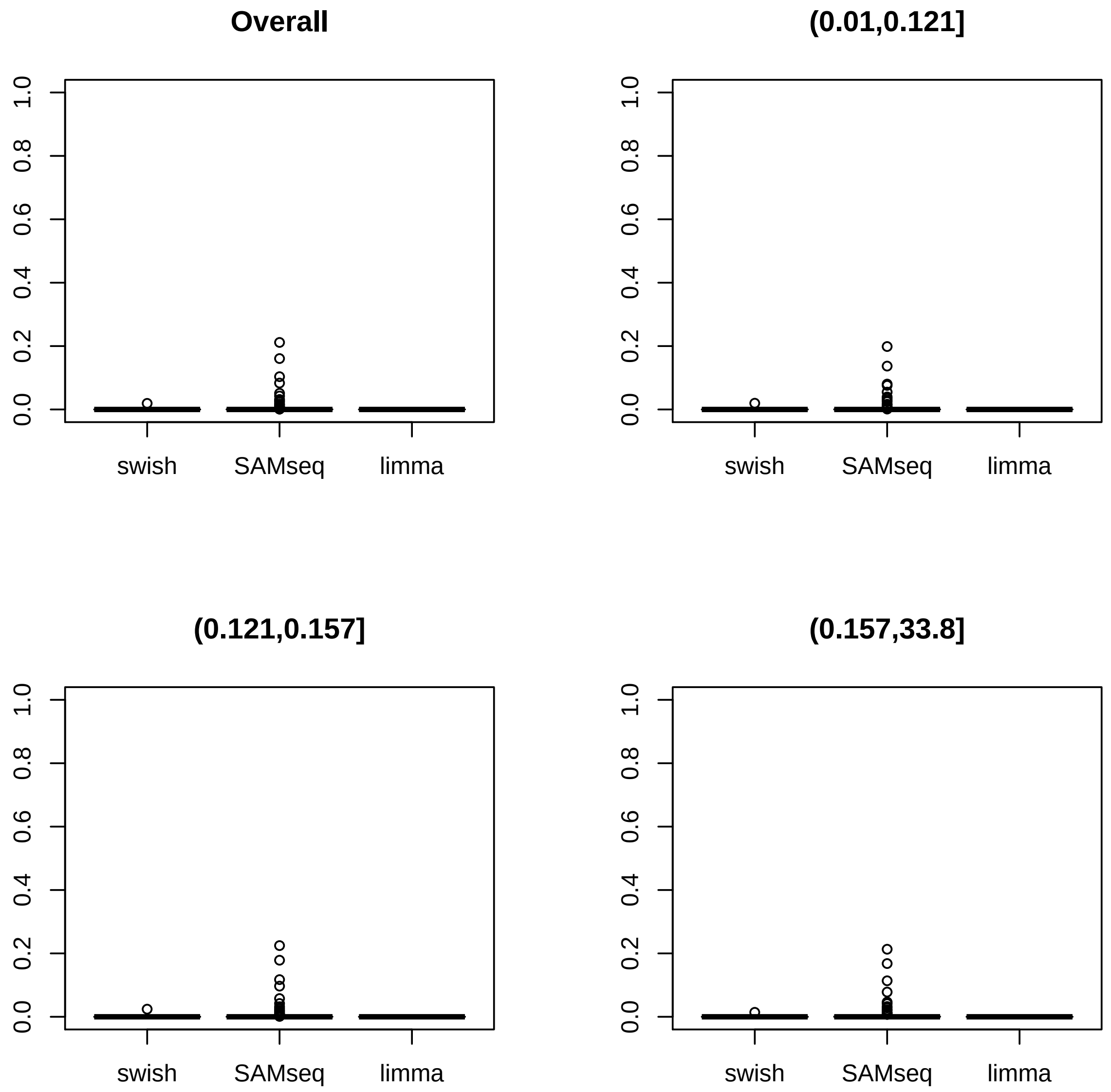


Supplementary Figure 28: False positive rate of *swish*, *SAMseq*, and *limma* over 100 repititions for yeast data at sample size 20. Panels provide overall FPR and FPR for features divided into thirds by InfRV.

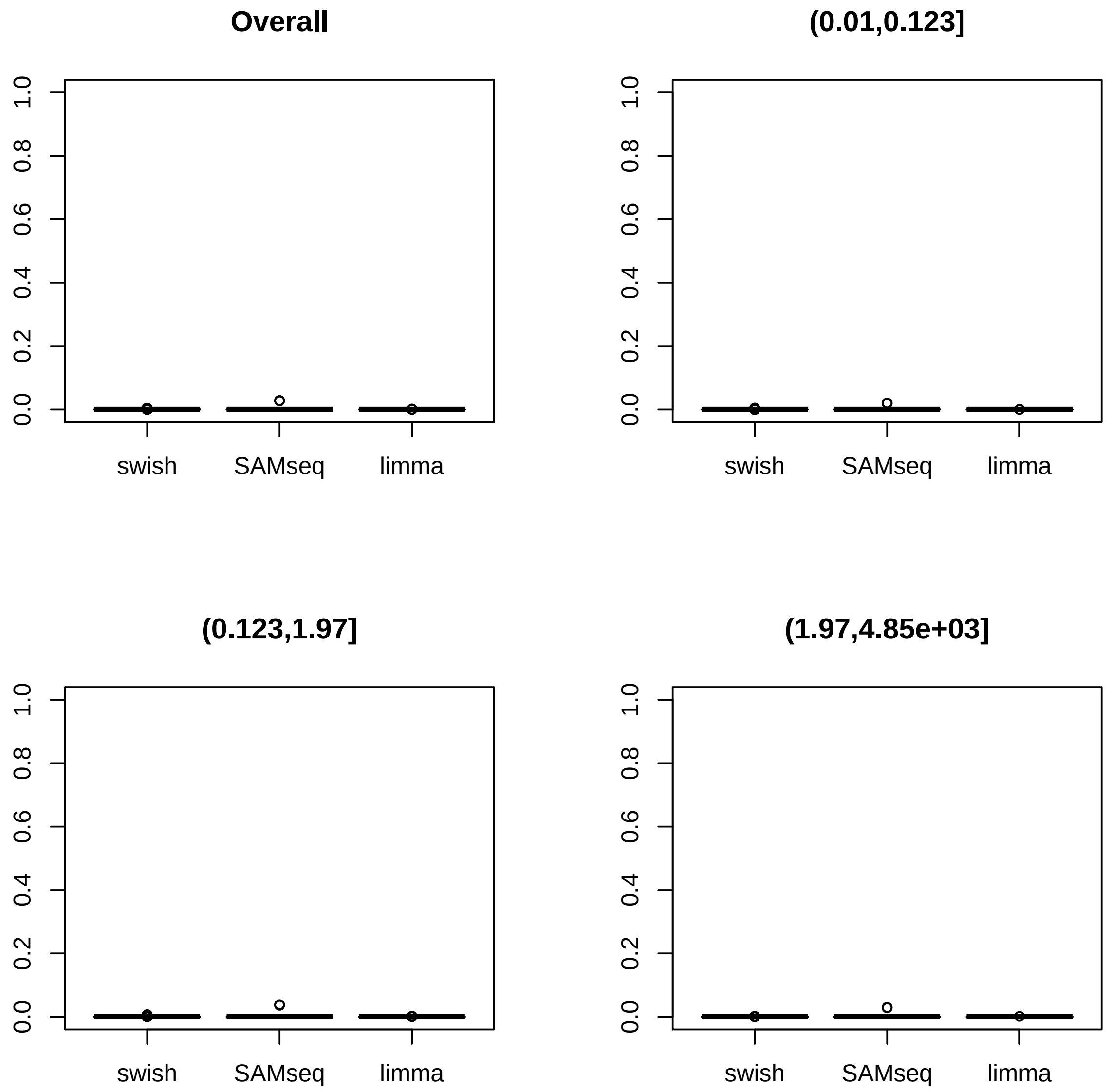


Supplementary Figure 29: False positive rate of *swish*, *SAMseq*, and *limma* over 100 repititions for transcript-level *Arabidopsis* data at sample size 5. Panels provide overall FPR and FPR for features divided into thirds by InfRV.

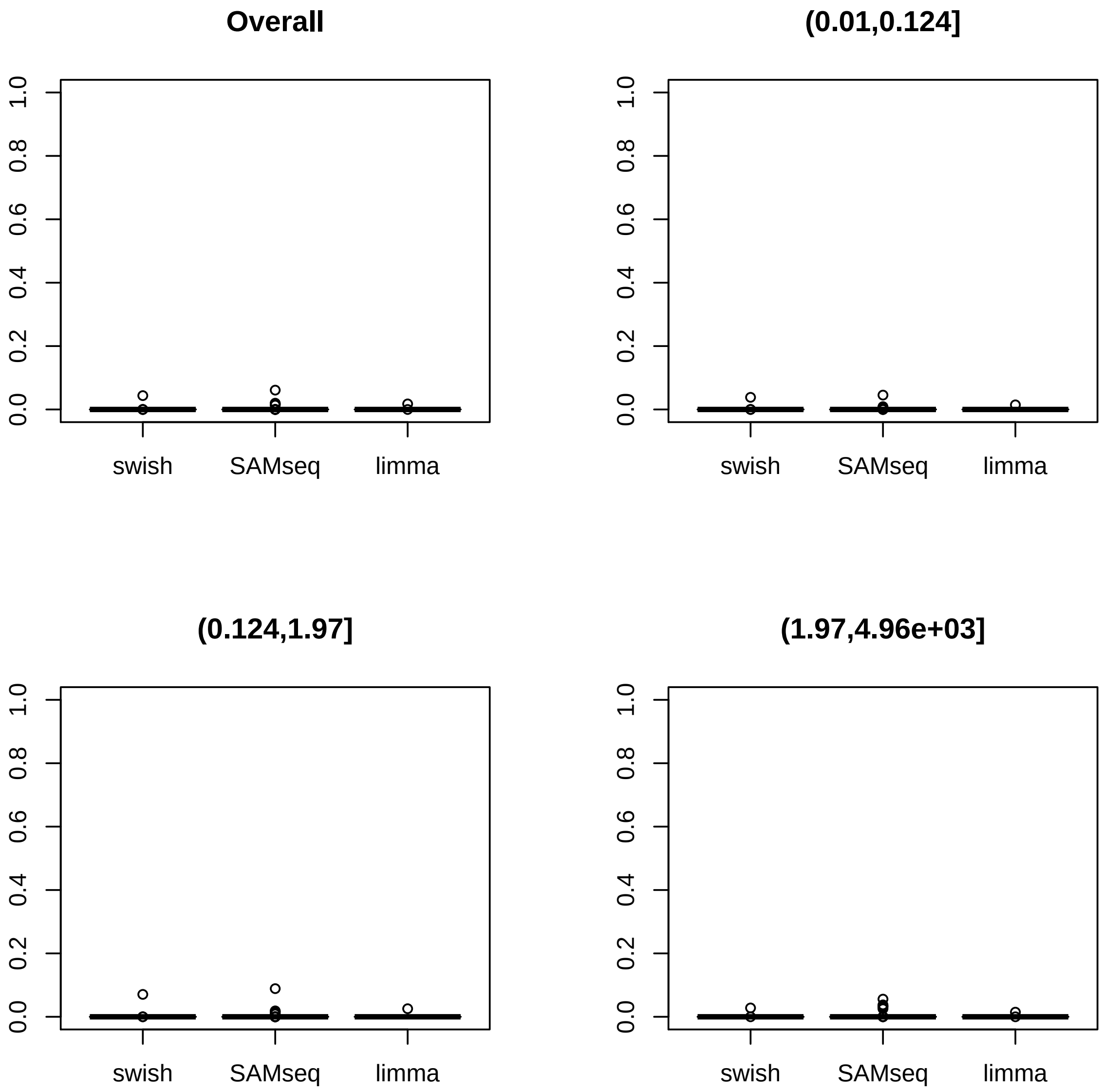


Supplementary Figure 30: False positive rate of *swish*, *SAMseq*, and *limma* over 100 repititions for transcript-level *Arabidopsis* data at sample size 8. Panels provide overall FPR and FPR for features divided into thirds by InfRV.

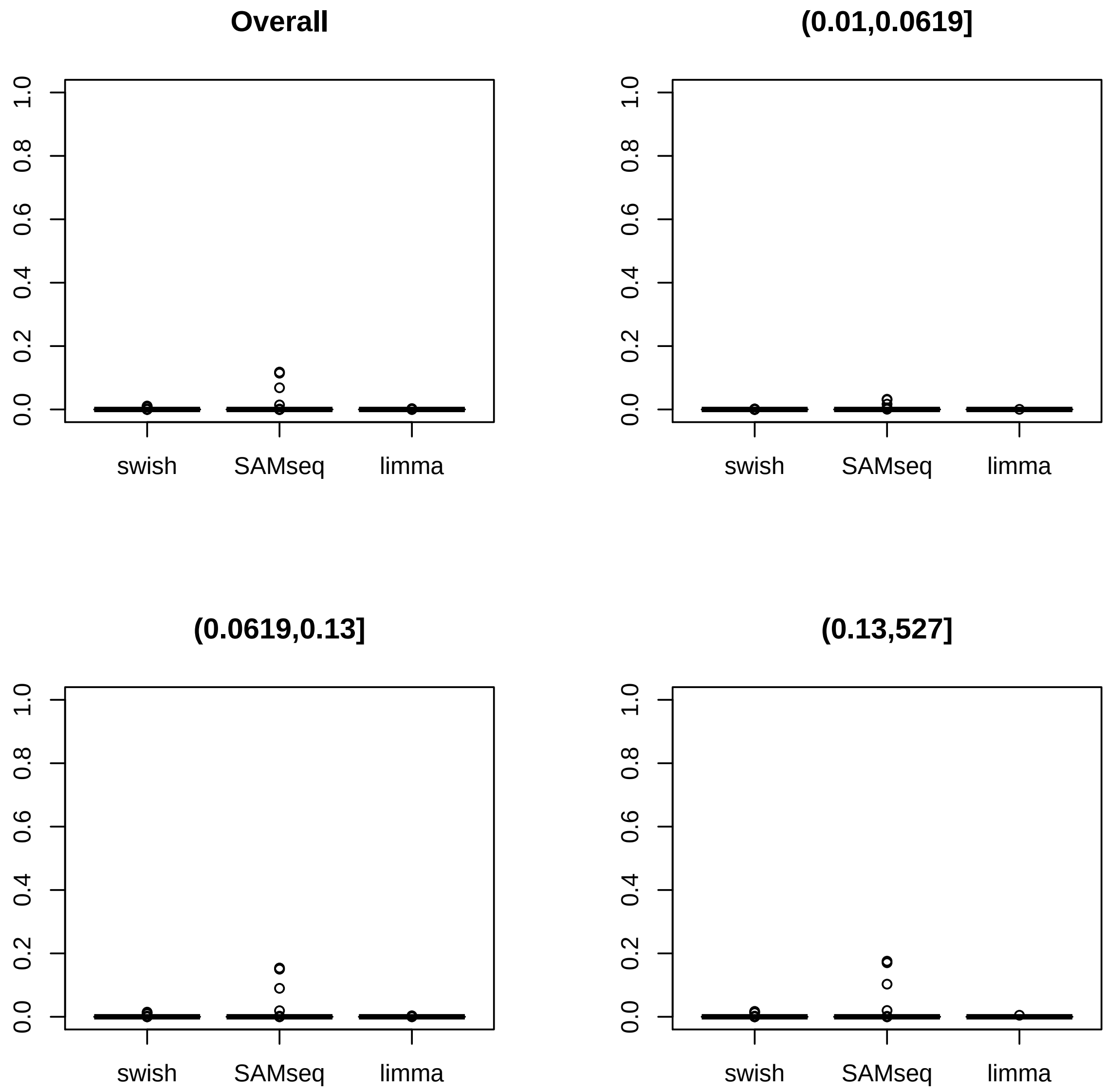


Supplementary Figure 31: False positive rate of *swish*, *SAMseq*, and *limma* over 100 repititions for gene-level *Arabidopsis* data at sample size 5. Panels provide overall FPR and FPR for features divided into thirds by InfRV.

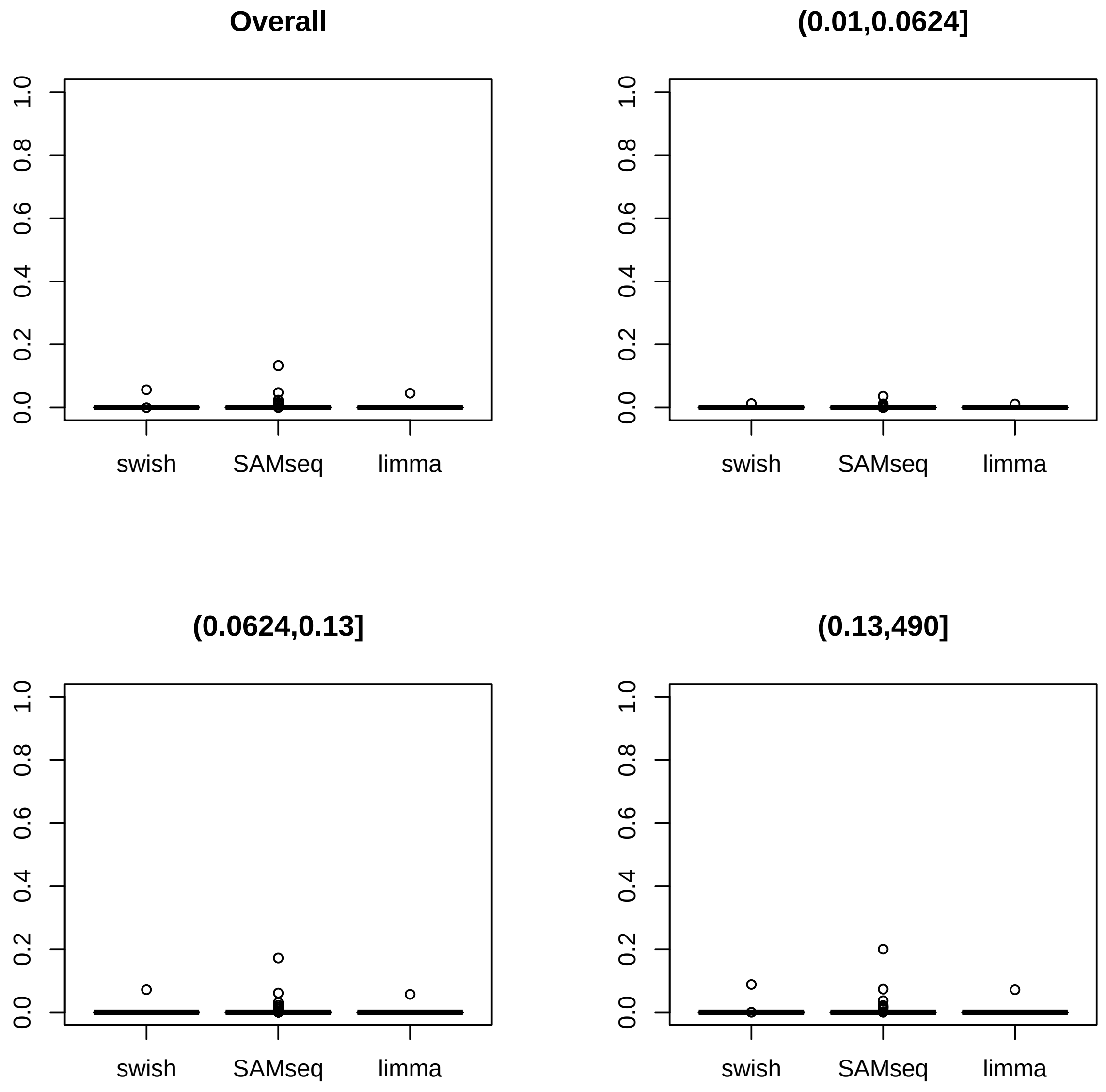


Supplementary Figure 32: False positive rate of *swish*, *SAMseq*, and *limma* over 100 repititions for gene-level *Arabidopsis* data at sample size 8. Panels provide overall FPR and FPR for features divided into thirds by InfRV.

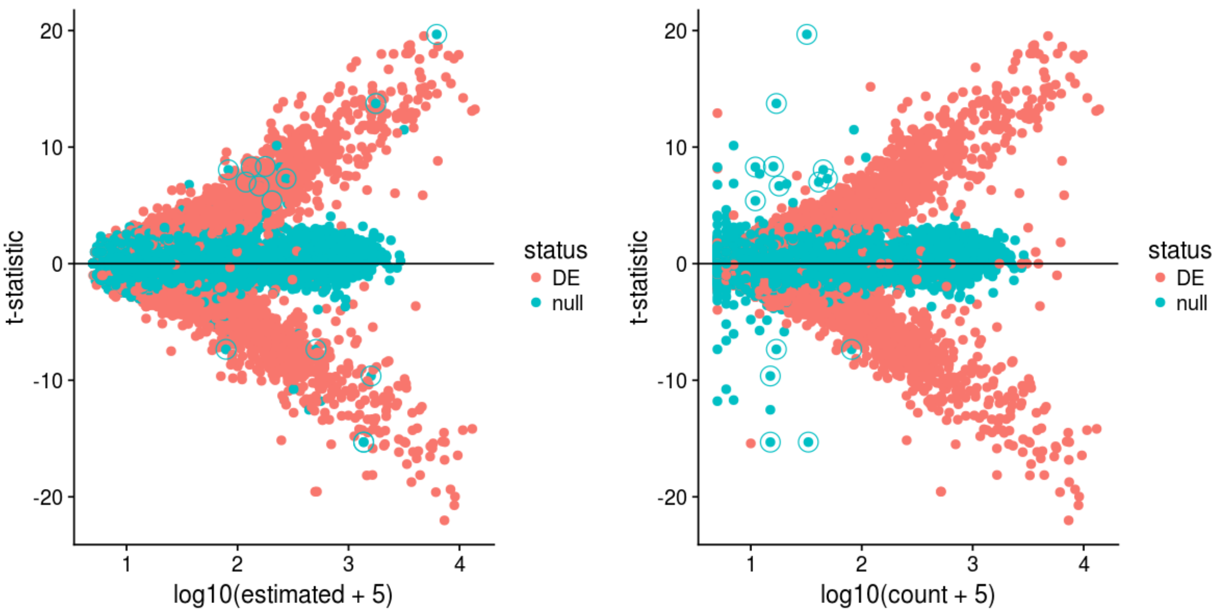


Supplementary Figure 33: MA-plot of *splatter-polyester* simulation of 40 cells, with t-statistic (y-axis) plotted across sum of counts per gene (x-axis). t-statistics were computed on the log of counts for visualization only and plotted over the $\log_{10}$ of total counts, whether estimated (left panel) or true counts (right panel). Of the null genes (blue), a number are revealed in the right panel as having large t-statistic despite being null genes. These genes contain regions of sequence homology with other, true DE genes, such that the null genes are assigned some non-zero proportion of the expression. 14 genes are highlighted with open circles in which the total count $> 5$, $|t| > 5$, and InfRV $> .2$. These are further examined in the following Supplementary Figures.

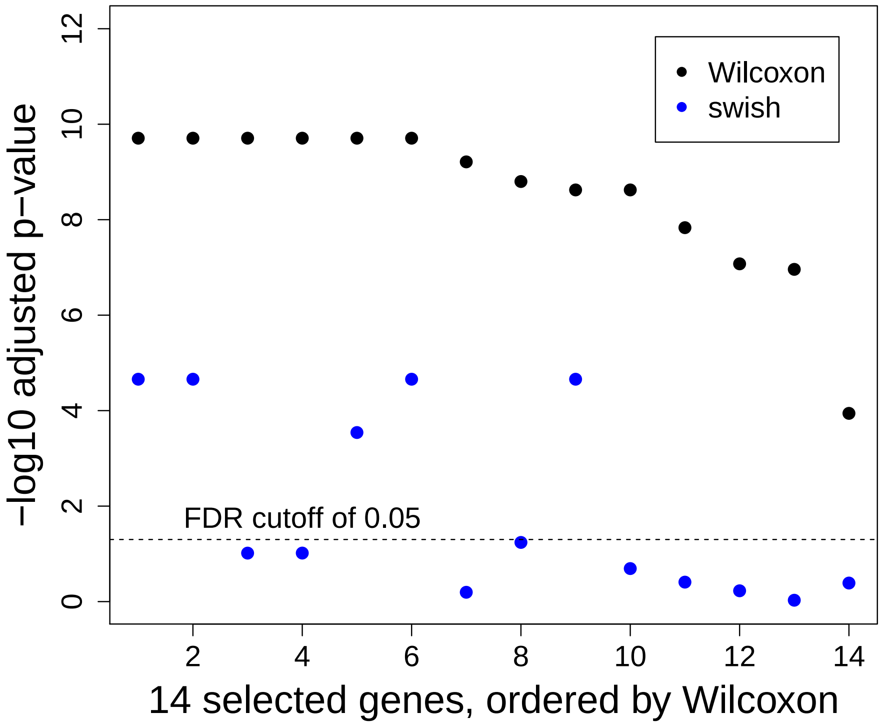


Supplementary Figure 34: Adjusted p-values for Wilcoxon test and *Swish*, for 14 genes from the *splatter-polyester* simulation, highlighted in the previous plot. These 14 null genes were chosen for having total count $> 5$, $|t| > 5$, and InfRV $> .2$. For all of these genes, *Swish* assigns a much higher q-value (less significant, and lower on the $-\log_{10}$ scale), and for 9 of the 14, the genes have such a high q-value that they are not included in a target 5% FDR gene set. These 9 false positive genes are not found significant by *Swish* due to their high inferential uncertainty, as seen in the following Supplementary Figure.

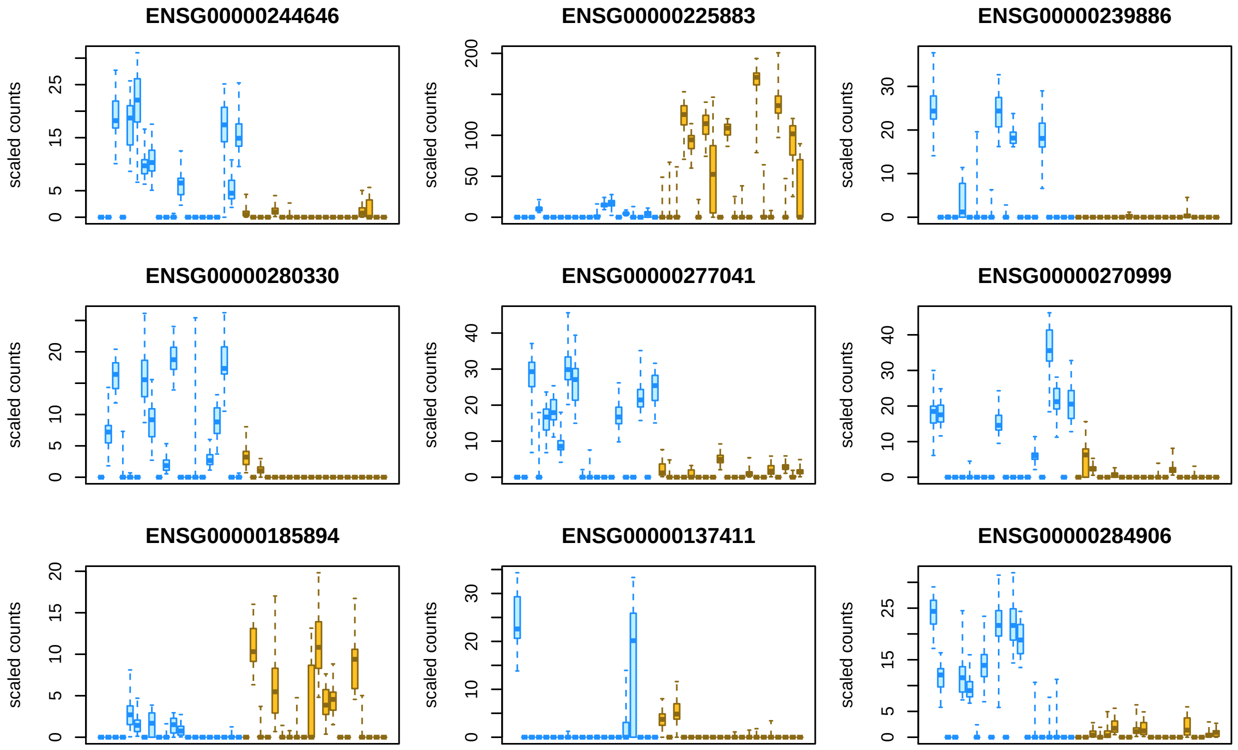


Supplementary Figure 35: Plots of inferential replicates for all 40 cells in the *splatter*-*polyester* simulation, for 9 null genes highlighted in the previous Supplementary Figure. These null genes were included in a 5% FDR set by Wilcoxon test followed by Benjamini-Hochberg p-value adjustment, but not by *Swish* q-value. The high expression in one group that can be seen in these plots is spurious, due to sequence homology with other differentially expressed genes in the simulation. *Swish* does not find these genes to be differentially expressed due to the high inferential uncertainty, as can be seen in the boxplots of inferential replicates per sample. Color and horizontal position indicates the 40 cells divided evenly into two groups of 20 cells each. The y-axis indicates scaled counts using *DESeq2* size factor estimation with `type="poscounts"`.

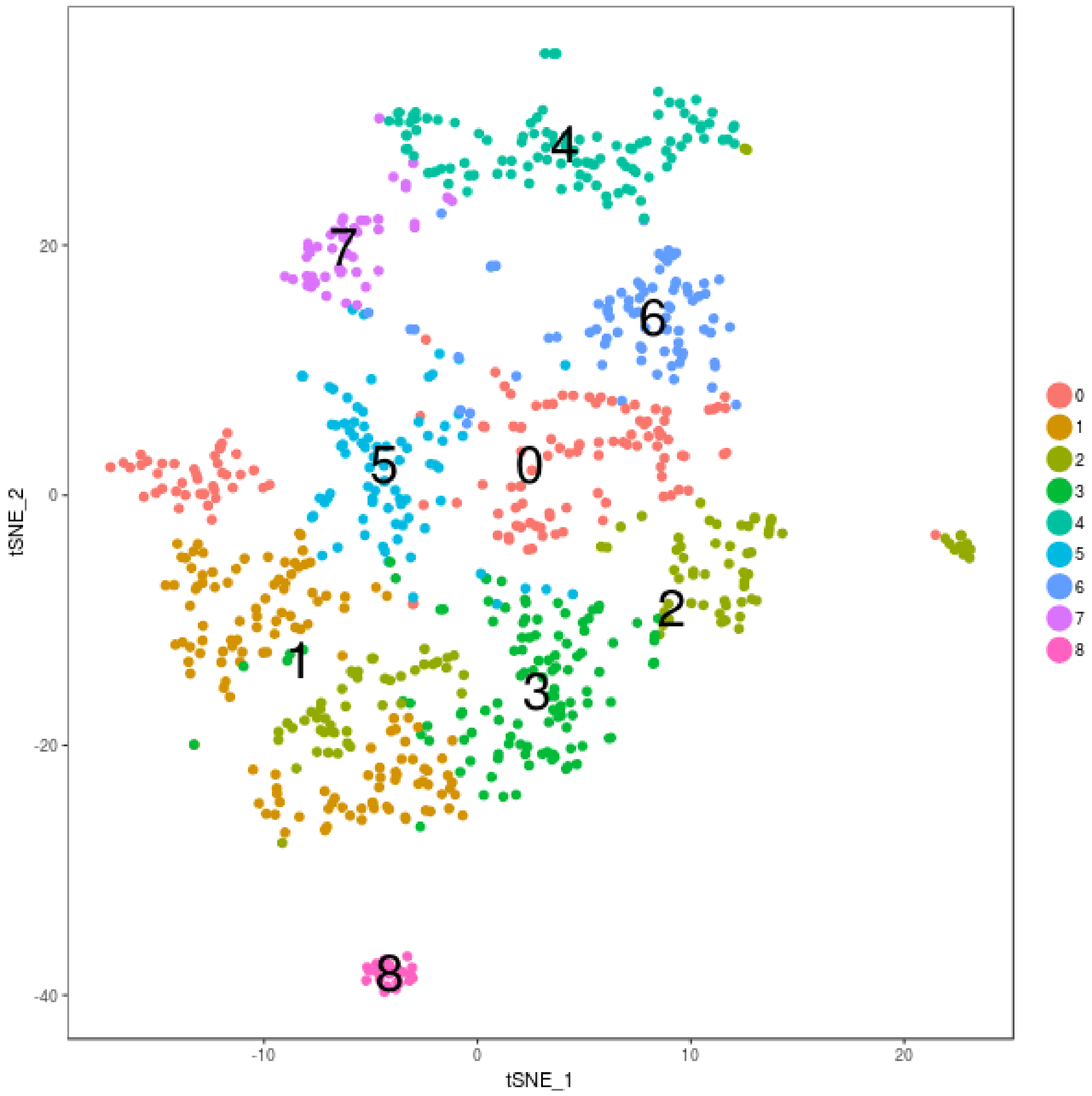


Supplementary Figure 36: The two-dimensional t-Distributed Stochastic Neighbor Embedding (tSNE) plot of the clustering results from *Seurat* for the developing mouse brain dataset. Each point represents a cell. Cells were cluster by *Seurat*'s graph-based method as described in the Methods, in the space of top PCs of log transformed, covariate adjusted data. Cells in the same cluster can be seen to localize together in the tSNE plot. Clusters are labeled by color and by the overlaid number.

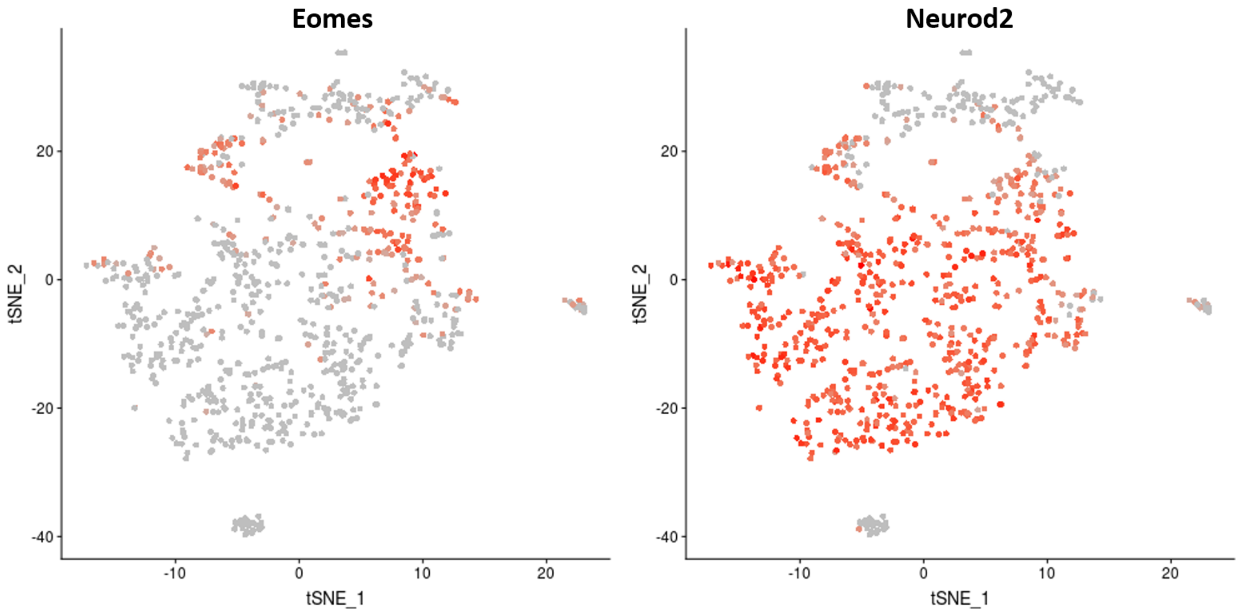


Supplementary Figure 37: The expression of two marker genes for the developing mouse brain, shown in the same tSNE plot as Supplementary Figure 36. Expression of *Eomes* (left) and *Neurod2* (right) per cell are visualized with red color. Grey dots indicate cells that do not have the gene expressed. *Eomes* is a marker for neural progenitor cells, and shows expression in clusters in the top of the figure. *Neurod2* is a marker for differentiation of neurons.

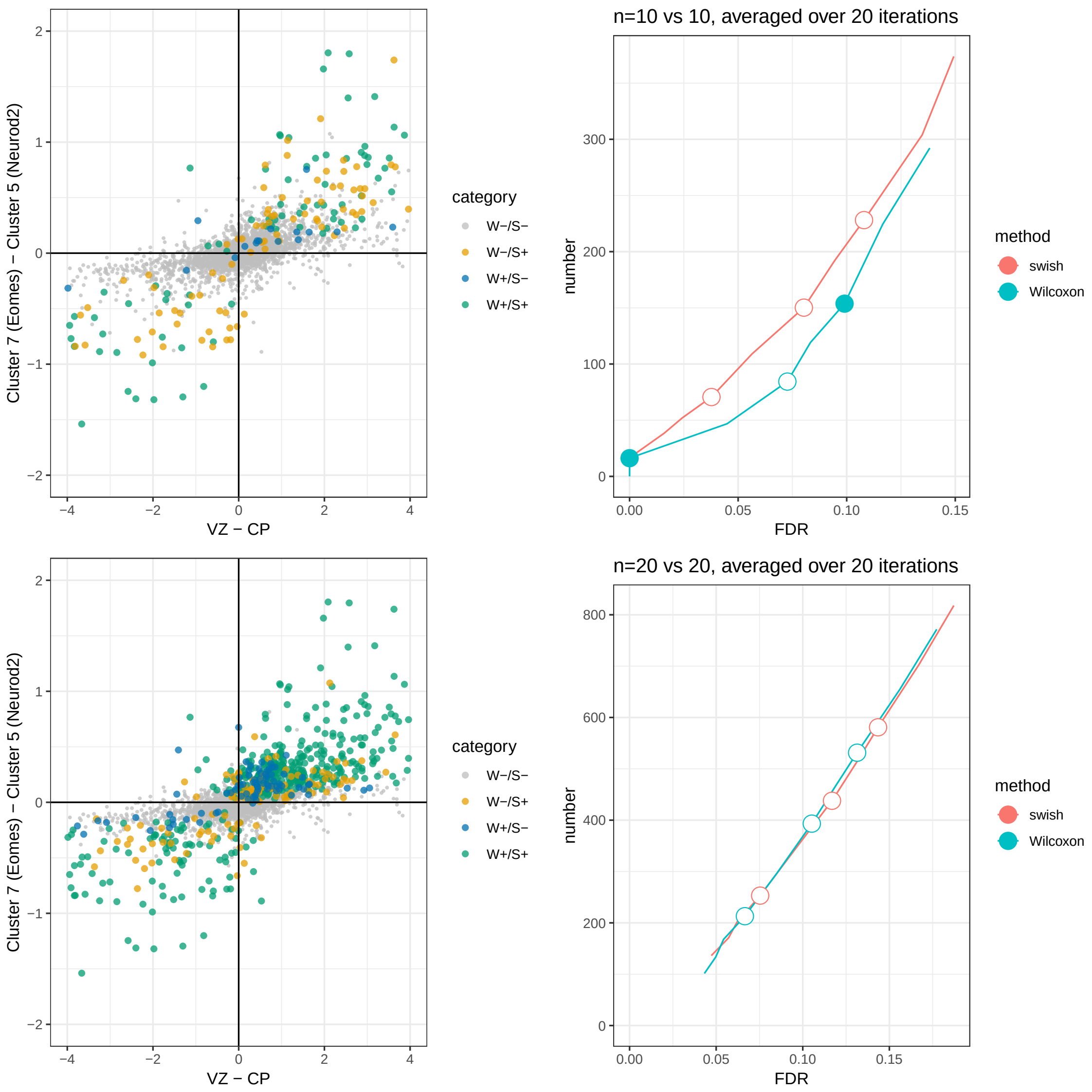


Supplementary Figure 38: Comparison of *Swish* and Wilcoxon test on the developing mouse brain dataset, with bulk RNA-seq as pseudo-gold-standard. The left panels show the log2 fold change from a comparison of cluster 7 and cluster 5 in the scRNA-seq dataset over the log2 fold change from a bulk RNA-seq dataset of the ventricular zone (VZ) and the cortical plate (CP). The Pearson correlation of the two sets of log2 fold changes is 0.61. The colors indicate those genes called by *Swish* only (yellow), Wilcoxon only (blue) or both methods (green). The right panels depict the number of genes detected by *Swish* and by the Wilcoxon test (y-axis) over the observed FDR (x-axis), with 1%, 5%, and 10% targets represented by circles (filled if the target FDR is achieved). The top row represents a comparison of 10 cells vs 10 cells and the bottom row represents a comparison of 20 cells vs 20 cells, drawn at random from the two clusters. The left panels give the colors from a single iteration, while the right panels give the average performance over 20 iterations of random subsets. A gene was declared a false positive if it was not detected as differentially expressed by *DESeq2* in the bulk RNA-seq data, or if the sign of the log2 fold change was inconsistent.

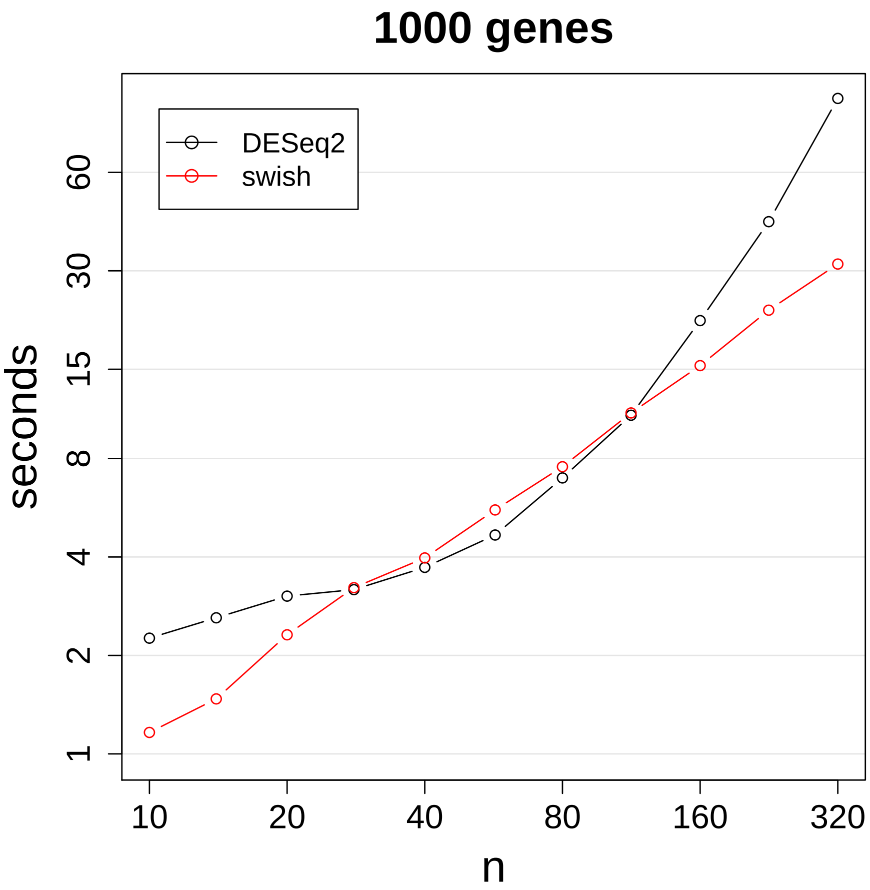


Supplementary Figure 39: Timing of *DESeq2* and *Swish* on the log-log scale, elapsed time over total sample size. *DESeq2* and *Swish* were run on 1,000 simulated genes with varying number of total samples (n) in a two group analysis, with 20 inferential replicates. *Swish* is faster than *DESeq2* for small sample sizes (total $n \leq 20$) and scales better, surpassing *DESeq2* at total sample size of 160, where *Swish* takes 15 seconds per 1,000 genes. While not shown here, both methods scale linearly with number of features, i.e. number of genes or transcripts.

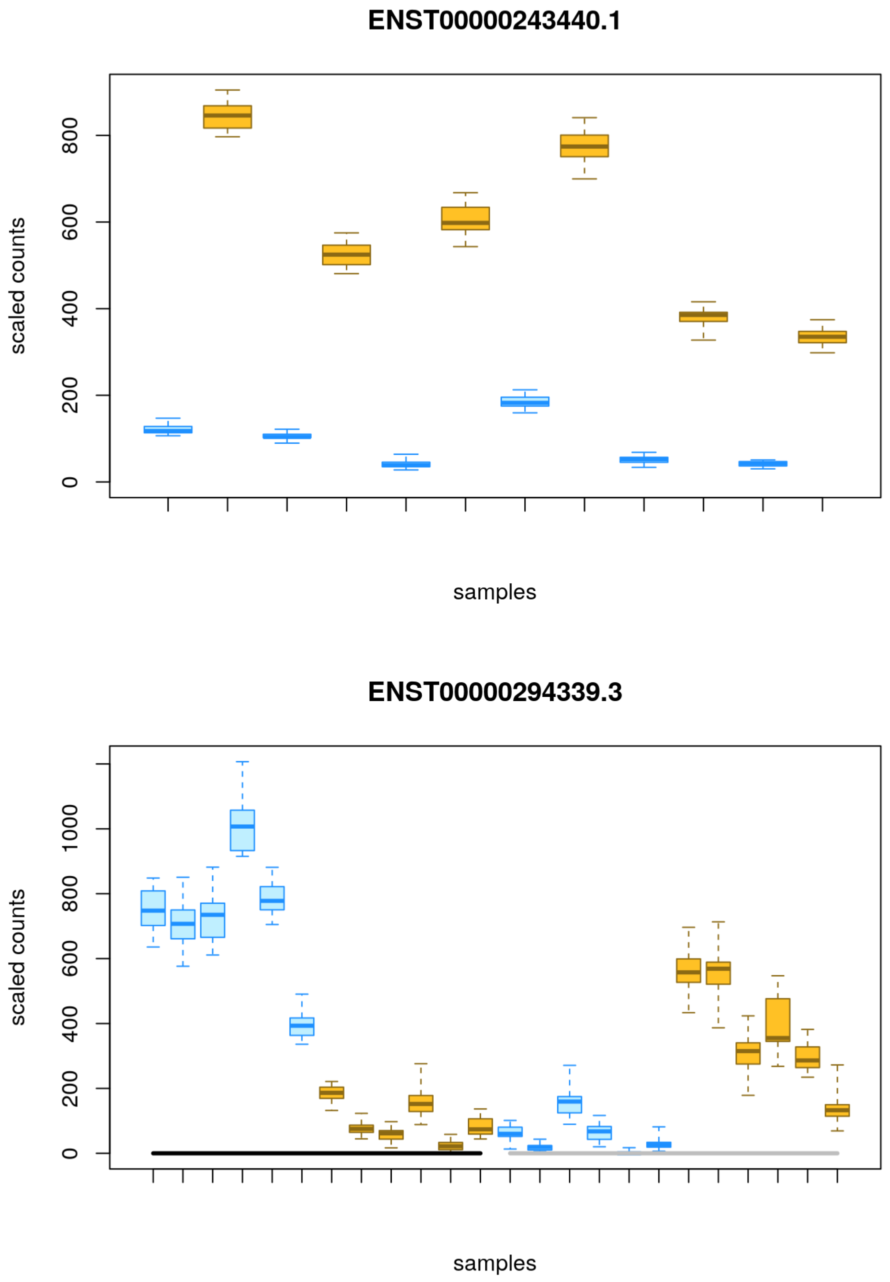


Supplementary Figure 40: Example inferential replicates plot from the `fishpond` software vignette on Bioconductor. Shown are transcripts identified as DTE in a paired two group comparison (top) or in a test of interaction between treatment with IFN-$\gamma$ and infection with *Salmonella* (bottom, black and grey groups denoting infection status). The paired two-group analysis and paired test for differences in treatment across groups are described in the Methods. The experimental data is a subset of data from Alasoo, *et al.* (2018) DOI: 10.1038/s41588-018-0046-7, where human macrophages were treated with immune stimulation via IFG-$\gamma$, infection with *Salmonella*, both treatments combined, or control. Each box in the boxplot denotes a single sample, where the box shows the spread of the inferential replicates. Matched samples from the same donor are consecutive. The blue boxes are control samples and the orange boxes are stimulated with IFN-$\gamma$.
